# IL-1β promotes MPN disease initiation by favoring early clonal expansion of *JAK2*-mutant hematopoietic stem cells

**DOI:** 10.1101/2023.05.21.541610

**Authors:** Shivam Rai, Yang Zhang, Elodie Grockowiak, Quentin Kimmerlin, Nils Hansen, Cedric B. Stoll, Marc Usart, Hui Hao-Shen, Michael S. Bader, Jakob R. Passweg, Stefan Dirnhofer, Christopher J. Farady, Timm Schroeder, Simón Méndez-Ferrer, Radek C. Skoda

## Abstract

*JAK2*-V617F is the most frequent somatic mutation causing myeloproliferative neoplasm (MPN). However, *JAK2*-V617F can also be found in healthy individuals with clonal hematopoiesis of indeterminate potential (CHIP) with a frequency much higher than the prevalence of MPN. The factors controlling the conversion of *JAK2*-V617F CHIP to MPN are largely unknown. We hypothesized that IL-1β mediated inflammation is one of the factors that favors this progression. We examined mono- or oligoclonal evolution of MPN by performing bone marrow transplantations at limiting dilutions with only 1-3 *JAK2*-mutant HSCs per recipient. Genetic loss of *IL-1β* in *JAK2*-mutant hematopoietic cells or inhibition by a neutralizing anti-IL-1β antibody restricted the early clonal expansion of these *JAK2*-mutant HSCs resulting in a reduced frequency of a CHIP-like state and a lower rate of conversion to MPN. The MPN disease-promoting effects of IL-1β were associated with damage to sympathetic innervation leading to loss of nestin-positive mesenchymal stromal cells and required the presence of *IL-1R1* on bone marrow stromal cells. The anti-IL-1β antibody protected these mesenchymal stromal cells from IL-1β mediated damage and limited the expansion of the *JAK2*-mutant clone. Our results identify IL-1β as a potential therapeutic target for preventing the transition from *JAK2*-V617F CHIP to MPN.

**Brief summary:** In a mouse model of oligo-clonal myeloproliferative neoplasm (MPN), IL-1β produced by *JAK2*-mutant cells favored expansion of sub-clinical *JAK2*-V617F clones and initiation of MPN disease.

## Introduction

Myeloproliferative neoplasms (MPNs) are clonal disorders of the hematopoietic stem cell (HSC) in most cases caused by activating mutations in *JAK2, CALR* or *MPL* that increase the proliferation of erythroid, megakaryocytic and myeloid lineages (1-7). MPNs can be subclassified as polycythemia vera (PV), essential thrombocythemia (ET) and primary myelofibrosis (PMF) (8, 9). *JAK2*-V617F is the most frequent driver gene mutation in patients with MPN, but *JAK2*-V617F can also be detected as “clonal hematopoiesis of indeterminate potential” (CHIP) in healthy individuals, with a frequency much higher than the incidence of MPN (10-13). Thus, the acquisition of *JAK2*-V617F is not the rate-limiting step for developing MPN, and there are factors that can suppress or promote the expansion of the *JAK2* mutated clone and thereby control the transition from CHIP to MPN.

Inflammation is a hallmark of advanced MPN (14). Chronic immune stimulation in patients with a history of infectious or autoimmune diseases was associated with increased risk of myeloid malignancies including MPN (15-17). Therefore, we hypothesized that inflammation might be one of the factors promoting the transition from *JAK2*-V617F positive CHIP to MPN. IL-1β, a pleiotropic cytokine with diverse innate and adaptive immune functions is a master regulator of inflammatory state (18, 19), and has been implicated in promoting several hematological malignancies (20, 21). Previous studies have shown that IL-1β is favoring progression to myelofibrosis at advanced stages of MPN (22, 23). In this study, we examined the role of IL-1β in the very early stages of MPN initiation by *JAK2*-V617F using competitive bone marrow (BM) transplantations at high dilutions (1:100), with very few (1-3) long-term hematopoietic stem cells (LT-HSCs). We found that IL-1β secreted by *JAK2*-mutant hematopoietic cells promotes the early expansion of the *JAK2*-V617F clone, whereas inhibition of IL-1β reduces MPN disease initiation by preventing IL-1β-induced destruction of sympathetic nerve fibers and thereby preserving nestin-positive mesenchymal stromal cells (MSCs).

## Results

### Loss of *IL-1β* from *JAK2*-mutant hematopoietic cells reduces the rate of MPN disease initiation

To test the hypothesis that IL-1β favors MPN disease initiation by promoting the early expansion of the *JAK2*-V617F clone, we performed competitive BM transplantations into lethally irradiated wildtype recipient mice. As BM donors, we used triple transgenic *SclCre^ER^;JAK2-*V617F;*UBC-GFP* (*VF;GFP*) mice that allow tracking of the transplanted BM cells by detection of GFP. These *VF;GFP* mice were further crossed to obtain quadruple transgenic *VF;IL-1β^-/-^;GFP* mice. To activate the *VF* transgene, these mice were injected with tamoxifen. After developing MPN phenotype, these mice were used as BM donors (Figure 1). To mimic the monoclonal origin of MPN from a *JAK2*-mutant HSC, we used competitive transplantations of unfractionated BM cells mixed at 1:100 ratio with *IL-1β^-/-^* BM competitor cells (Figure 1A), resulting in transplantation of only 1-3 *JAK2*-mutant LT-HSCs per recipient (24). Within 36 weeks, about 60% of recipients transplanted with *VF;GFP* and *IL-1β^-/-^* BM competitor cells developed MPN phenotype characterized by elevated hemoglobin and/or platelet counts and splenomegaly (Figure 1A). Mice that developed MPN phenotype also showed increased IL-1β levels in plasma and BM, while mice without MPN phenotype displayed very low IL-1β levels. Very similar results were obtained in experiments when wildtype (*WT*) BM competitor cells were used instead of *IL-1β^-/-^* BM (Supplemental Figure S1), indicating that *WT* competitor cells did not contribute to the increased IL-1β production.

**Figure 1.**
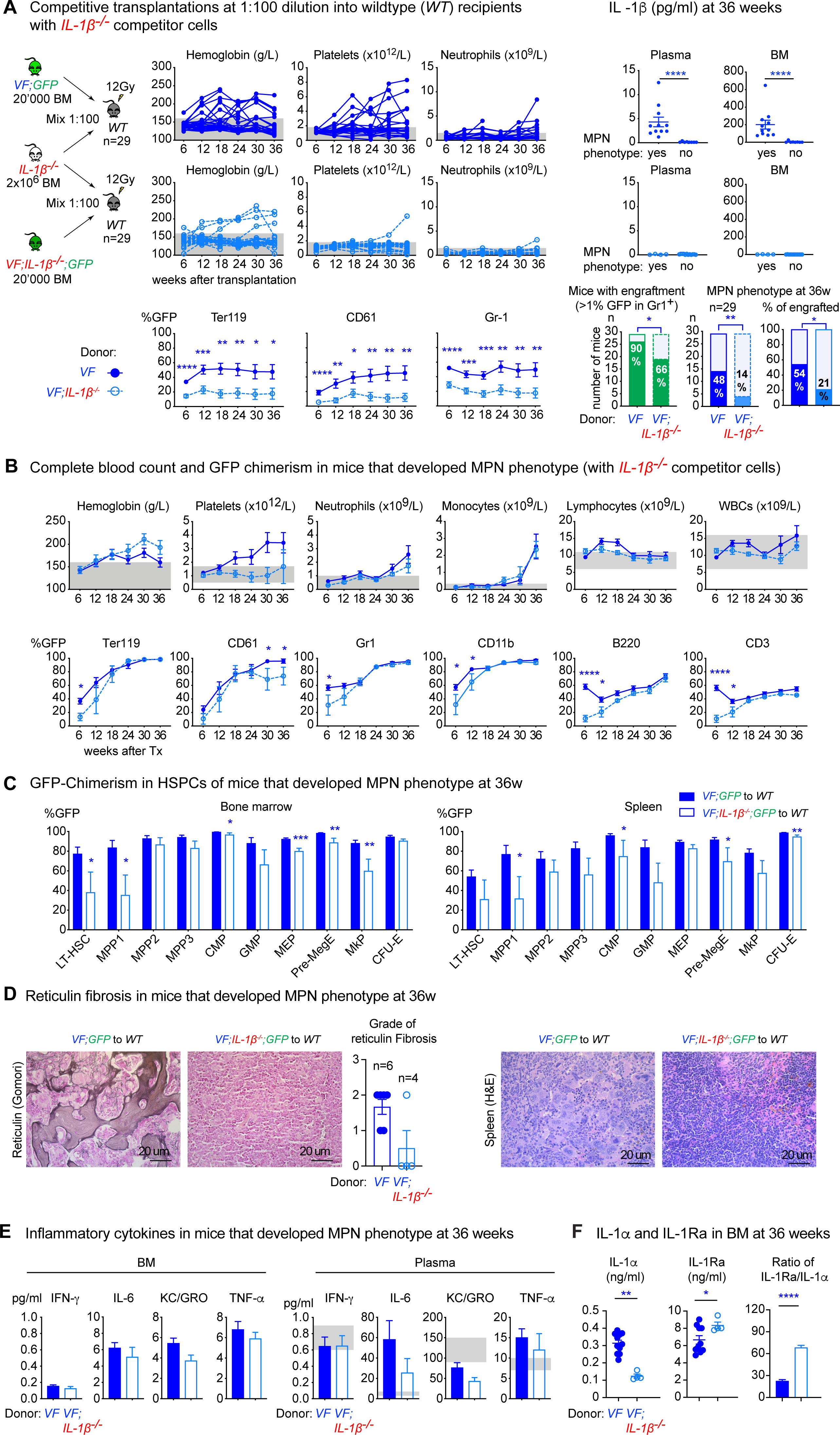
Loss of *IL-1β* from *JAK2*-mutant hematopoietic cells reduces MPN disease initiation. **(A)** Schematic drawing of the experimental setup for competitive transplantation at 1:100 dilution. Bone marrow (BM) from *VF;GFP* or *VF;IL-1β^-/-^;GFP* donor mice was mixed with a 100-fold excess of BM competitor cells from a *IL-1β^-/-^* donor. The time course of blood counts from individual mice that received BM from *VF;GFP* (upper panel) or *VF;IL-1β^-/-^;GFP* donors (middle panel), and the GFP chimerism in peripheral blood (lower panel) are shown. Multiple t tests were performed for statistical analyses. IL-1β protein levels in plasma and BM lavage (1 femur and 1 tibia) of mice with or without MPN phenotype is shown (right panel). Non-parametric Mann-Whitney two-tailed t test was performed for statistical comparisons. Bar graphs (bottom right) show the percentages of mice that showed engraftment defined as GFP-chimerism >1% at 18 weeks after transplantation and the percentages of mice that developed MPN phenotype (elevated hemoglobin and/or platelet counts). p values in lower panel were computed using Fisher’s exact test. **(B)** Time course of mean blood counts and GFP chimerism in peripheral blood of *WT* mice transplanted with BM from *VF;GFP* or *VF;IL-1β-/-;GFP* and *IL-1β^-/-^* competitor cells that developed MPN phenotype during 36-weeks follow-up. Multiple t tests were performed for statistical analyses. **(C)** GFP-chimerism in hematopoietic stem and progenitor cells (HSPCs) at 36 weeks after transplantation in BM (left) and spleen (right) of *WT* mice transplanted with BM from *VF;GFP* or *VF;IL-1β-/-;GFP* and *IL-1β^-/-^* competitor cells that developed MPN phenotype. Multiple t tests were performed for statistical analyses. **(D)** Representative images of reticulin fibrosis staining in BM (left panel) and H&E staining in spleen (right panel) of mice showed MPN phenotype at 36 weeks after transplantation. Histological grade of reticulin fibrosis in BM is shown in a bar graph (right). **(E)** Levels of Inflammatory cytokines in BM lavage (1femur and 1 tibia) and plasma of mice that displayed MPN phenotype at 36 weeks after transplantation. **(F)** Levels of IL-1 cytokines in BM. Grey shaded area represents normal range. All data are presented as mean ± SEM. *P < .05; **P < .01; ***P < .001; ****P < .0001. See also Supplemental Figure S1 and S2.

We have previously shown that *VF* and *VF;IL-1β^-/-^* mice have comparable frequencies of HSCs in BM (22). When we transplanted BM cells from a *VF;IL-1β^-/-^;GFP* donor mixed with *IL-1β^-/-^* BM competitor cells at 1:100 ratio, we found that considerably fewer recipients developed MPN phenotype compared to mice that received *VF;GFP* BM and only one mouse showed thrombocytosis (Figure 1A, middle panel). Recipients without MPN phenotype did not show increased IL-1β levels in plasma or BM, suggesting that excess IL-1β was produced by the *JAK2-*V617F expressing cells. Recipients transplanted with *VF;GFP* BM showed higher GFP-chimerism (Figure 1A, lower panel), higher percentage of engraftment (90% versus 66%), defined as GFP-chimerism >1% in peripheral blood Gr1^+^ granulocytes, and a higher percentage of MPN disease initiation (48% versus 12%) than mice transplanted with *VF;IL-1β^-/-^;GFP* BM (Figure 1A, lower panel).

When only recipients that developed MPN phenotype are considered, blood counts and GFP-chimerism in the peripheral blood were similar in mice transplanted with *VF;GFP* versus *VF;IL-1β^-/-^;GFP* BM Figure 1B). However, platelets as well as GFP chimerism in CD61 cells in peripheral blood were slightly higher in recipients of *VF;GFP* BM. Furthermore, in mice that developed MPN phenotype, loss of *IL-1β* from *JAK2*-mutant cells resulted in reduced GFP-chimerism in hematopoietic stem and progenitor cells (HSPCs) in BM and spleen (Figure 1C), less reticulin fibrosis and osteosclerosis in BM, and partial restoration of splenic architecture (Figure 1D). Inflammatory cytokines were largely unchanged, except for a trend towards lower IL-6 levels (Figure 1E). Loss of *IL-1β* in *VF* donor cells resulted in reduced levels of IL-1α in the BM of mice that developed MPN (Figure 1F). Interestingly, loss of *IL-1β* in *VF* donor cells also increased the levels of IL-1 receptor antagonist (IL-1Ra) in these mice. As a result, the ratio of IL-1Ra to IL-1α was increased in recipients transplanted with *VF;IL-1β^-/-^*BM that showed MPN phenotype, correlating with the less severe MPN phenotype and less myelofibrosis, compared with recipients transplanted with *VF* BM.

Finally, transplantations of *VF* BM with *IL-1β^-/-^* competitor cells into *IL-1β^-/-^* recipients resulted in elevated IL-1β levels (Supplemental Figure S2A, upper panel), similar to transplantations into wildtype recipients, demonstrating that the source of overproduction of IL-1β were hematopoietic cells that express *JAK2*-V6167F. Unexpectedly, *IL-1β^-/-^* recipients transplanted with *VF;IL-1β^-/-^* BM showed frequencies of engraftment and MPN disease initiation that were higher than anticipated and almost equal to *IL-1β^-/-^* recipients transplanted with *VF* BM (Supplemental Figure S2A, middle panel). The GFP chimerism was reduced when all recipient mice were considered (Supplemental Figure S2A, lower panel), but when only mice that developed MPN were compared, few alterations were noted only at some of the time points (Supplemental Figure S2B). Nevertheless, *IL-1β^-/-^* recipients transplanted with *VF;IL-1β^-/-^* BM showed a trend towards reduced reticulin fibrosis, reduced GFP-chimerism in HSPCs and partial restoration of splenic architecture (Supplemental Figure S2C-D). A trend towards reduction of IL-6 and KC/GRO levels was observed in BM and plasma (Supplemental Figure S2E).

The reasons why engraftment and MPN disease initiation increased when IL-1β was absent in both donors and recipients is currently unknown. No increase in IL-1Ra or in the ratio between IL-1α and IL-1Ra was found in *IL-1β^-/-^* recipients transplanted with *VF;IL-1β^-/-^* BM (Supplemental Figure S2F), suggesting that lack of upregulation of IL-1RA might contribute to the more efficient MPN disease initiation. Complete loss of *IL-1β* partially prevented this outcome, possibly due to diminished anti-IL-1 inflammatory responses in the BM.

Overall, our data show that loss of *IL-1β* in hematopoietic cells reduced MPN initiation, suggesting that IL-1β produced by the *JAK2*-mutant cells is promoting efficient early expansion of the *JAK2-*V617F clone and optimal MPN disease initiation.

### *JAK2*-V617F mutant HSCs need IL-1β for optimal expansion of the HSC population with long-term repopulation capacity

To examine the effects of *IL-1β* loss on mutant stem cell function, we performed secondary transplantations into wildtype recipients (Figure 2). As donors, we selected 3 mice transplanted with *VF;GFP* and 3 mice transplanted with *VF;IL-1β^-/-^;GFP* BM that displayed MPN phenotype (Supplemental Figure S3). The BM from all 3 donors was pooled and used for non-competitive and competitive secondary transplantations (Figure 2). Loss of *IL-1β* from donor *VF* BM resulted in significantly reduced blood counts and lower GFP chimerism in peripheral blood, BM and spleen of secondary recipients, compared with mice that received *VF* donor cells with wildtype *IL-1β* (Figure 2A-B and Supplemental Figure S4A-B). By comparing the engraftment frequencies (defined as >1% in Gr1+ granulocytes at 18 weeks) between the competitive and non-competitive secondary transplantations, and using the “extreme limiting dilution analysis” algorithm (25), we estimated that functional HSCs were ∼5-fold reduced in donor mice that were initially transplanted with *VF;IL-1β^-/-^* BM compared to donors initially transplanted with *VF* BM (Figure 2C). Overall, the HSC pool in the primary recipient mice transplanted with 1-3 HSCs from *VF* donors expanded to a total of ∼ 300 functional HSCs after 36 weeks, whereas primary recipients transplanted with 1-3 HSCs from *VF;IL-1β^-/-^* donors expanded to a total of only ∼50 functional HSCs during the same time. Taken together, these results indicate that *JAK2*-V617F mutant HSCs require IL-1β for optimal expansion of the HSC population with long-term repopulation capacity.

**Figure 2.**
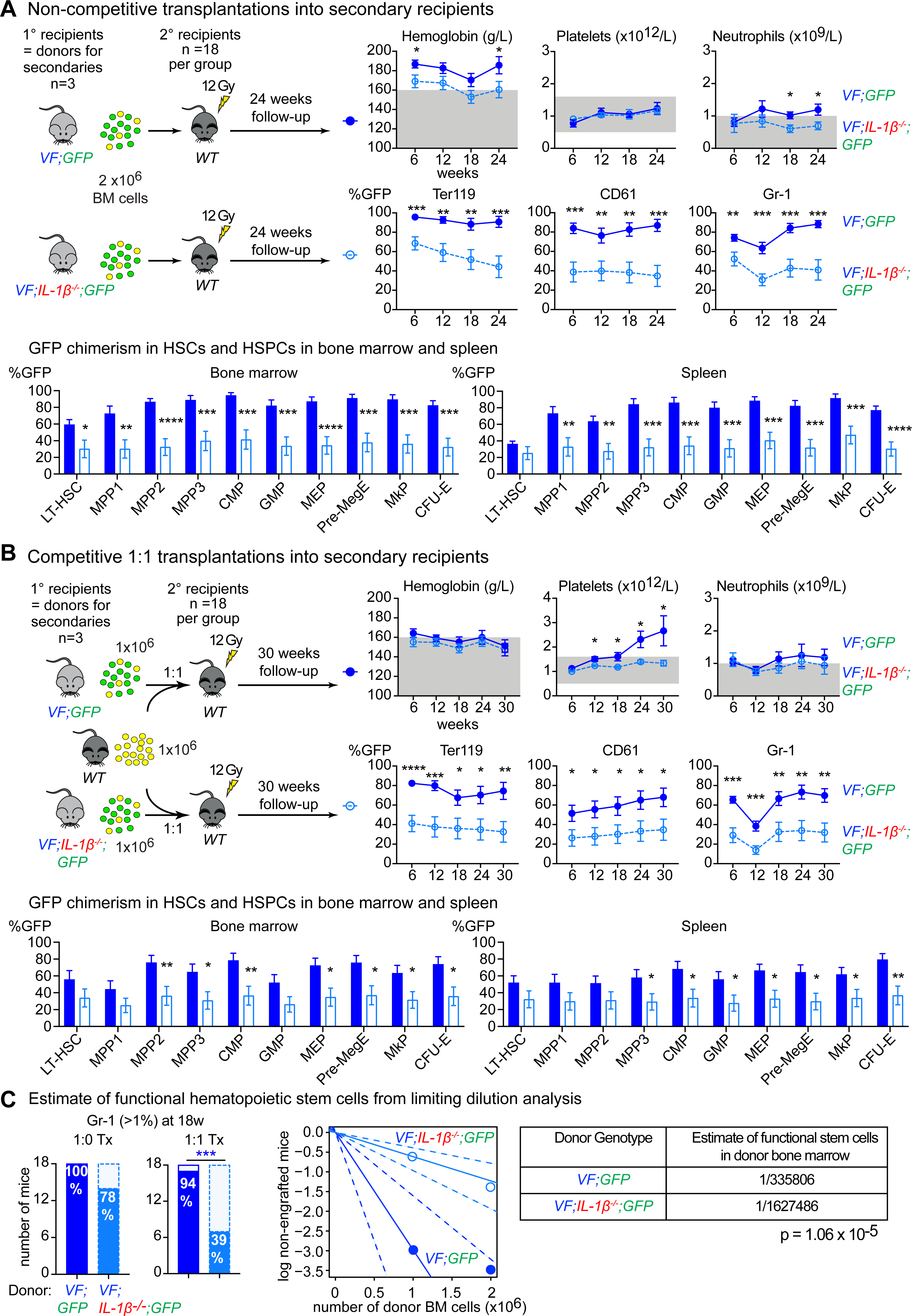
*JAK2*-V617F HSCs need IL-1β for long-term stem cell function. **(A)** Schematic drawing of non-competitive (1:0) transplantations. Primary recipients of *VF;GFP* or *VF;IL-1β^-/-^;GFP* bone marrow (BM) were sacrificed at 36 weeks after transplantation and their BM cells (2×10^6^) were transplanted into *WT* recipients (n=18 per group). The time course of blood counts from mice that received BM from *VF;GFP* (solid symbols) or *VF;IL-1β^-/-^;GFP* donors (open symbols), and their GFP chimerism in peripheral blood (lower panel) are shown. Analysis of GFP chimerism in hematopoietic stem cells and progenitors from BM and spleen is shown below. Multiple t tests were performed for statistical analyses. **(B)** Schematic drawing of the identical experiment as above, in which BM competitor cells from a *WT* mouse was mixed 1;1 ratio for the transplantations into secondary *WT* recipients. Annotations as in (A). Multiple t tests were performed for statistical analyses. **(C)** Bar graphs show the percentages of mice that showed engraftment defined as GFP-chimerism >1% at 18 weeks after non-competitive (left) and competitive 1:1 transplantation (right). Middle panel shows the log of non-engrafted mice (GFP-chimerism in Gr1<1% at 18-weeks) versus the number of donor BM cells transplanted in each group. Estimated frequency of functional stem cells in BM was calculated using extreme limiting dilution analysis.(25) Solid dark blue line represents the estimated frequency of HSC in mice transplanted with BM from *VF;GFP* donors. Solid light blue line represents the estimated frequency of HSC in mice transplanted with BM from *VF;IL-1β^-/-^;GFP* donors. Dotted lines show 95% confidence interval. Grey shaded areas represent the normal range. All data are presented as mean ± SEM. *P < .05; **P < .01; ***P < .001; ****P < .0001. See also Supplemental Figure S3 and S4.

### MPN initiation by *JAK2*-V617F is favored by signaling through the IL-1R1 in both hematopoietic and non-hematopoietic cells

We next examined the role of *IL-1R1* in MPN disease initiation. We used BM cells from the same *VF;GFP* donor as in Figure 1, or a *VF;IL-1R1^-/-^;GFP* donor to perform competitive transplantations with a 100x excess of BM competitor cells from a *IL-1R1^-/-^* mouse into lethally irradiated *WT* recipients (Figure 3A). Engraftment was observed in 97% of recipients and 55% developed MPN phenotype characterized by elevated hemoglobin and/or platelet counts, with significantly elevated IL-1β levels compared to mice without MPN phenotype (Figure 3A, upper panel). In mice transplanted with BM from *VF;IL-1R1^-/-^;GFP*, engraftment was 100% but only 27% of recipients developed MPN phenotype.

**Figure 3.**
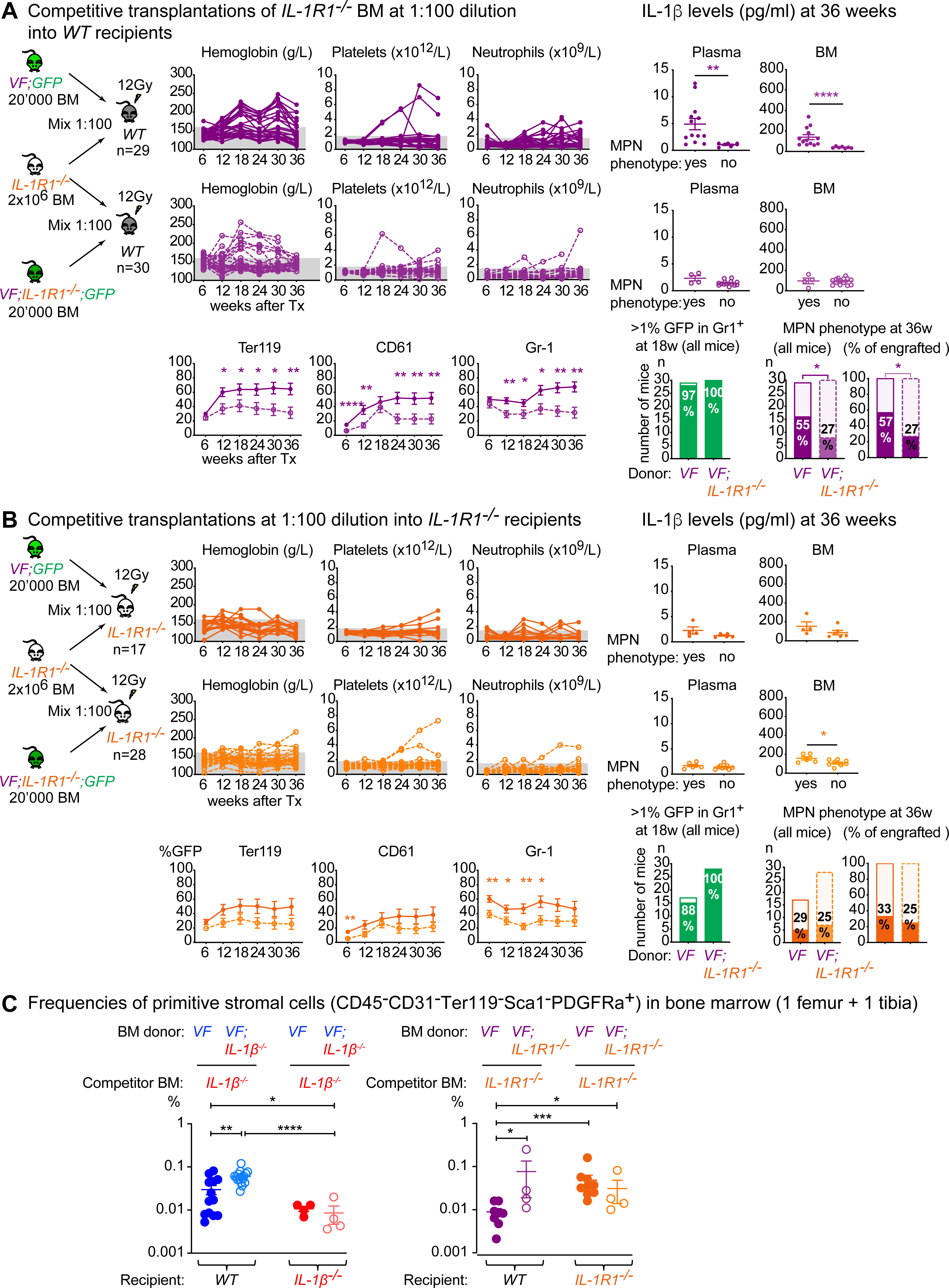
*JAK2*-mutant cells need IL-1R1 expression on both hematopoietic and non-hematopoietic cells for optimal MPN initiation. **(A)** Schematic drawing of the experimental setup for competitive transplantation at 1:100 dilution. Bone marrow (BM) from *VF;GFP* or *VF;IL-1R1^-/-^;GFP* donor mice was mixed with a 100-fold excess of BM competitor cells from a *IL-1R1^-/-^* donor. The time course of blood counts from individual mice that received BM from *VF;GFP* (upper panel) or *VF;IL-1R1^-/-^;GFP* donors (middle panel), and the GFP chimerism in peripheral blood (lower panel) are shown. Multiple t tests were performed for statistical analyses. IL-1β protein levels in plasma and BM lavage (1 femur and 1 tibia) of mice with or without MPN phenotype is shown (right panel). Non-parametric Mann-Whitney two-tailed t test was performed for statistical comparisons. Bar graphs (bottom right) show the percentages of mice that showed engraftment defined as GFP-chimerism >1% at 18 weeks after transplantation and the percentages of mice that developed MPN phenotype (elevated hemoglobin and/or platelet counts). p values in lower panel were computed using Fisher’s exact test. **(B)** Schematic drawing of the identical experiment as above, in which *IL-1R1^-/-^* mice were used as the recipients instead of *WT* mice. Annotations as in (A). **(C)** Frequency of primitive stromal cells (CD45^-^CD31^-^Ter119^-^Sca1^-^PDGFRα^+^) in the BM (1 femur and 1 tibia) of mice from Figure 1A (left panel) and Figure 3A and 3B (right panel) is shown. Non-parametric Mann-Whitney two-tailed t test was performed for statistical comparisons. Grey shaded area represents normal range. All data are presented as mean ± SEM. *P < .05; **P < .01; ***P < .001; ****P < .0001. See also Supplemental Figure S5 and S6.

GFP-chimerism in peripheral blood was higher in mice transplanted with *VF;GFP* BM than in recipients of *VF;IL-1R1^-/-^;GFP* BM, when all mice that showed engraftment were considered (Figure 3A), whereas GFP-chimerism was similar when only mice that developed MPN phenotype were compared (Supplemental Figure S5A). However, loss of *IL-1R1* from *JAK2*-mutant cells was associated with reduced GFP chimerism in HSPCs in BM and spleen (Supplemental Figure S5B). Recipients of *VF;IL-1R1^-/-^;GFP* BM displayed low IL-1β levels in plasma and BM and no differences were noted between mice with or without MPN phenotype (Figure 3A, middle panel). Nevertheless, these IL-1β levels were higher compared with mice transplanted with *VF;IL-1β^-/-^*BM (Figure 1A). While *VF;IL-1β^-/-^*BM cells are unable to produce IL-1β, *VF;IL-1R1^-/-^* BM cells can still produce IL-1β, although at lower levels than *VF* BM cells, due to loss of the positive IL-1β-feedback mediated by the IL-1R1. Loss of *IL-1R1* in *JAK2*-mutant cells also reduced reticulin fibrosis and osteosclerosis in BM and partially corrected the splenic architecture in mice that developed MPN phenotype (Supplemental Figure S5C).

Similar transplantations were performed into recipients deficient for *IL-1R1^-/-^* (Figure 3B). Engraftment was as efficient as in transplantations into wildtype recipients, but MPN disease initiation in *IL-1R1^-/-^* deficient recipients transplanted with *VF;GFP* BM was reduced to 29% (Figure 3B), compared to 55% in *WT* recipients (Figure 3A). In mice that developed MPN phenotype, blood counts did not change, but GFP-chimerism in peripheral blood was slightly reduced with complete loss of *IL-1R1* (Supplemental Figure S6A), while GFP-chimerism remained unchanged in HSPCs in BM and spleen (Supplemental Figure S6B). Compared to *WT* recipients, *IL-1R1^-/-^* recipients transplanted with *VF;GFP* showed reduced grade of reticulin fibrosis in BM, normalization of splenic architecture (Supplemental Figure S6C) and reduced levels of inflammatory cytokines (Supplemental Figure S6D). However, loss of *IL-1R1* from both donor and recipient did not affect reticulin fibrosis or the levels of inflammatory cytokines (Supplemental Figure S6C-D).

In all mice with engraftment, we examined the frequencies of primitive BM mesenchymal stromal cells (MSCs) (CD45-CD31-Ter119-Sca1-PDGFRα+) that are enriched in nestin-positive MSCs (26-28). Transplantation of *VF* BM into *WT* recipients resulted in lower numbers of MSCs compared with transplantation of *VF;IL-1β^-/-^* BM into *WT* recipients (Figure 3C), indicating that IL-1β derived from *VF* cells reduces the numbers of primitive BM MSCs. In contrast, transplantation of *VF* or *VF;IL-1R1^-/-^* hematopoietic cells into *IL-1R1^-/-^* recipient mice failed to reduce primitive BM MSCs, suggesting that *IL1R1* signaling within the primitive BM MSCs causes their own reduction.

Taken together, these results show that increased expression of *IL-1β* by the *JAK2*-mutant hematopoietic cells and the presence of *IL-1R1* on both the mutant hematopoietic cells and the wildtype non-hematopoietic cells are required for efficient MPN disease initiation and progression.

### Inhibiting inflammation can reduce MPN disease initiation in a mouse model of *JAK2*-V617F driven clonal hematopoiesis

Since reducing pro-inflammatory signaling by *IL-1β* knockout in *VF* donor cells or deletion of *IL-1R1* in the recipient cells both reduced MPN disease initiation in competitive transplantations at high dilution, we examined whether the same effect can be observed with pharmacologic inhibition of inflammation. We first tested the effects of aspirin, which was added to the drinking water on day 1 after transplantation (Supplemental Figure S7). We observed a slight reduction in the frequencies of engraftment (96% in the controls versus 70% in the aspirin treated mice), but there were no differences in the blood counts or frequencies of MPN disease initiation (Supplemental Figure S7). Next, we treated recipients of *VF;GFP* BM cells with anti-IL-1β antibody for 18 weeks, starting on day 1 after transplantation (Figure 4A). Anti-IL-1β antibody treatment reduced the percentage of mice with engraftment >1% and fewer mice developed MPN phenotype compared to mice treated with isotype control (Figure 4B and C). Control mice transplanted with BM from *WT:GFP* donor showed no differences in phenotype or percentages of engraftment between the anti-IL-1β antibody and isotype treatment groups (Supplemental Figure S8). IL-1β concentration in BM and plasma was massively reduced in mice treated with anti-IL-1β antibodies (Figure 4D). HSPCs from mice with MPN phenotype displayed high GFP-chimerism that did not change by anti-IL-1β antibody treatment, whereas HSPCs from mice with engraftment >1% but without MPN phenotype, resembling a state of CHIP, showed low GFP-chimerism and in some cases a decrease in chimerism was noted upon anti-IL-1β treatment (Figure 4E).

**Figure 4.**
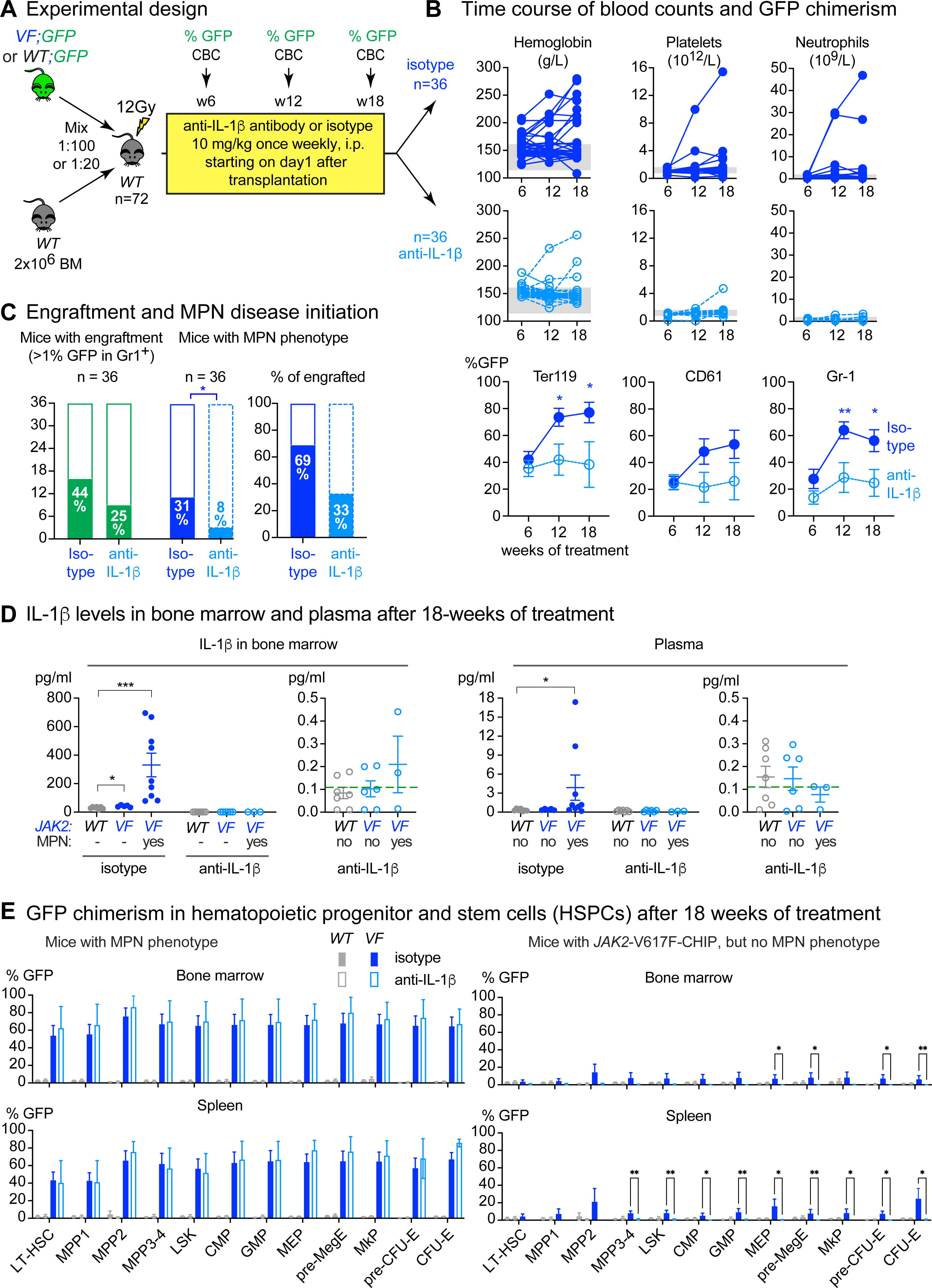
Anti-IL-1β antibody reduces MPN disease initiation in a mouse model of *JAK2*-V617F driven clonal hematopoiesis. **(A)** Schematic drawing of the experimental setup for competitive transplantation at 1:100 dilution. Bone marrow (BM) from a *VF;GFP* donor mouse was mixed with a 100-fold excess of BM competitor cells from a *WT* donor. Mice were randomized into two treatment arms one day after transplantation and treated with either isotype or anti-IL-1β antibody for 18 weeks. **(B)** The time course of blood counts from individual mice for the two treatment arms is shown (top and middle panel). Mean GFP (mutant cell) chimerism in peripheral blood erythroid (Ter119), megakaryocytic (CD61), granulocytic (Gr1) cells are shown in the bottom panel. Multiple t tests were performed for statistical analyses. **(C)** Bar graph showing the percentage of mice that showed engraftment defined as GFP-chimerism >1% at 18 weeks after transplantation and the percentage of mice that developed MPN phenotype (elevated hemoglobin and/or platelet counts). p value in right panel was computed using Fisher’s exact test. **(D)** Bar graph showing IL-1β protein levels in BM lavage (1 femur and 1 tibia) and plasma of mice with or without *JAK2*-V617F and/or MPN phenotype. Non-parametric Mann-Whitney two-tailed t test was performed for statistical comparisons. Green dashed line represents limit of detection. **(E)** Bar graphs in the left panel shows the mutant GFP chimerism within hematopoietic stem and progenitor cells in BM (upper panel) and Spleen (lower panel) of mice with *JAK2*-V617F and MPN phenotype at 18 weeks after transplantation. Bar graphs in the right panel shows the chimerism in mice with *JAK2*-V617F CHIP but no MPN phenotype. Multiple t tests were performed for statistical analyses. All data are presented as mean ± SEM. *P < .05; **P < .01; ***P < .001; ****P < .0001. See also Supplemental Figure S7 and S8.

Hematopoietic cells expressing *JAK2*-V617F were previously shown to cause loss of sympathetic nerve fibers and the Schwann cells that surround them, resulting in a reduction of nestin-positive BM MSCs, which participate in forming the niche for HSCs. The loss of these cells was associated with MPN disease manifestations (21). A reduction of primitive BM MSCs, sympathetic nerve fibers and Schwann cells was observed in all isotype treated mice (Figure 5A-C), including mice that showed engraftment, but did not develop MPN, resembling a state of CHIP. Treatment with the anti-IL-1β antibody prevented the reduction of primitive BM MSCs (Figure 5A) and using immunofluorescence staining on skulls, we found that anti-IL-1β antibodies also prevented the reduction of sympathetic nerve fibers and Schwann cells (Figure 5B and C). These protective effects of anti-IL-1β antibody resemble the protective effects observed in *IL-1R1* knockout recipients transplanted with *VF* BM cells (Figure 3C).

**Figure 5.**
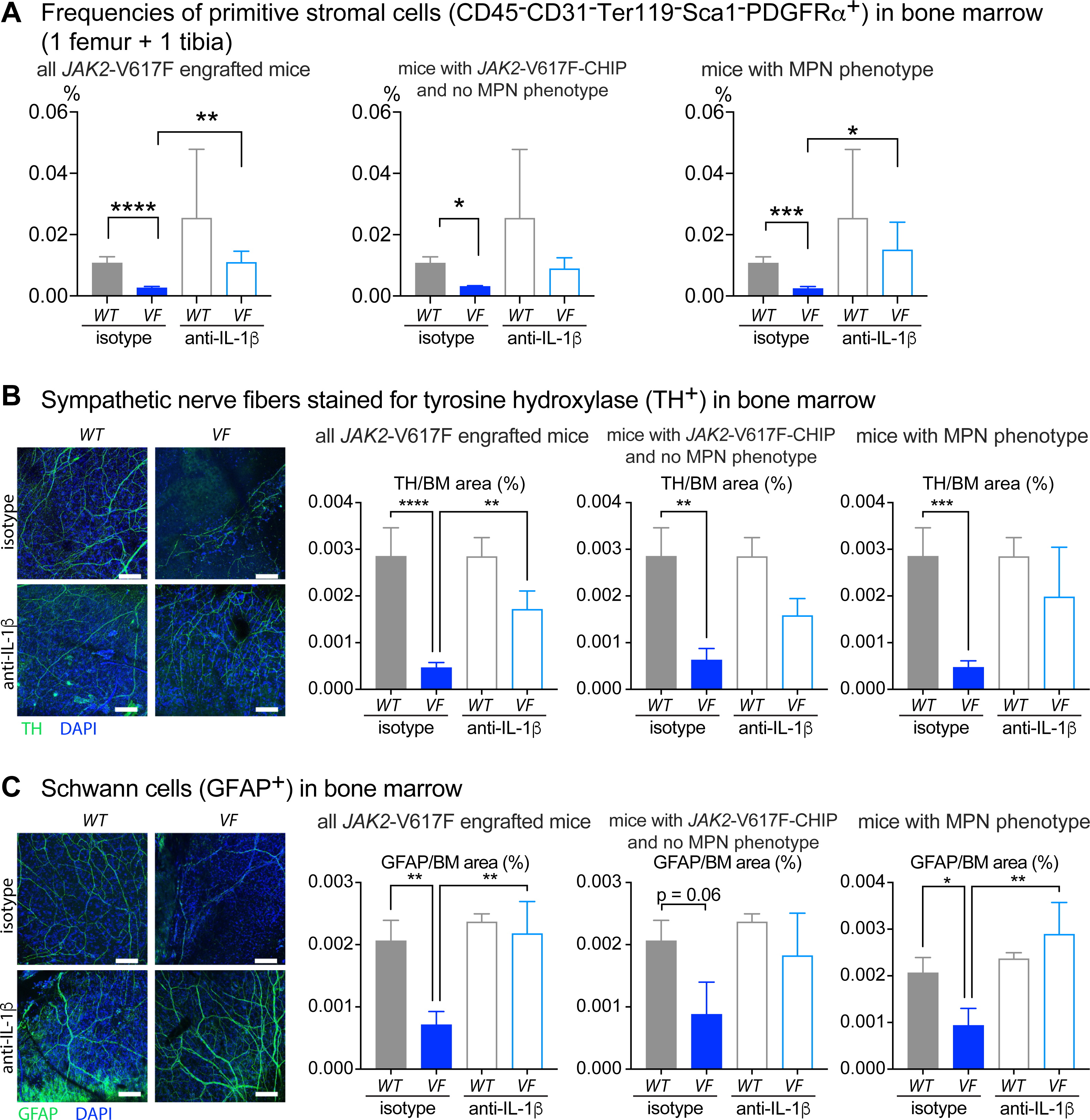
Inhibition of IL-1β preserves the HSC niche in the bone marrow. **(A)** Frequency of primitive stromal cells (CD45-CD31-Ter119-Sca1-PDGFRα+) in skull of mice transplanted with BM from *WT;GFP* or *VF;GFP* donor mouse after 18 weeks of treatment with isotype or anti-IL-1β antibody (from Figure 4). Bar graphs showing the frequencies of stromal cells in all engrafted mice (left panel), mice with *JAK2*-V617F-positive CHIP but no MPN phenotype (middle panel) and mice with MPN phenotype (right panel). Two-tailed unpaired t tests were performed for statistical comparisons. **(B)** Representative images of tyrosine hydroxylase (TH)-positive sympathetic nerve fibers (left panels) in skull bone marrow (BM) of mice transplanted with BM from *WT;GFP* or *VF;GFP* donor mouse after 18 weeks of treatment with isotype or anti-IL-1β antibody (from Figure 4). Bar graphs (right panels) showing the quantification of TH area in all engrafted mice (left panel), mice with *JAK2*-V617F-positive CHIP but no MPN phenotype (middle panel) and mice with MPN phenotype (right panel). One-way ANOVA with uncorrected Fisher’s LSD test was performed for statistical comparisons. **(C)** Representative images of glial fibrillary acidic protein (GFAP)-positive Schwann cells (left panels) in skull of mice transplanted with BM from *WT;GFP* or *VF;GFP* donor mouse after 18 weeks of treatment with isotype or anti-IL-1β antibody (from Figure 4). Bar graphs (right panels) showing the quantification of GFAP area in all engrafted mice (left panel), mice with *JAK2*-V617F-positive CHIP but no MPN phenotype (middle panel) and mice with MPN phenotype (right panel). One-way ANOVA with uncorrected Fisher’s LSD test was performed for statistical comparisons All data are presented as mean ± SEM. *P < .05; **P < .01; ***P < .001; ****P < .0001.

### Myelo-monocytic cells and megakaryocytes are main sources of IL-1β production in the bone marrow of *VF* mice

To analyze the distribution of IL-1β protein in bone marrow sections, we used in situ bone marrow 3D fluorescence imaging with Proximity Ligation Assay (PLA) that can visualize single IL-1β molecules (29-31). *WT* mice showed low densities of IL-1β PLA signals, whereas *VF* mice that developed MPN phenotype displayed the highest densities of PLA signals (Figure 6 and Supplemental Figure S9A). IL-1β PLA signals were enriched in close proximity of CD11b+ myeloid cells in mice of all genotypes, and these gradients were reduced in VF mice that were treated with anti-IL-1β antibody.

**Figure 6.**
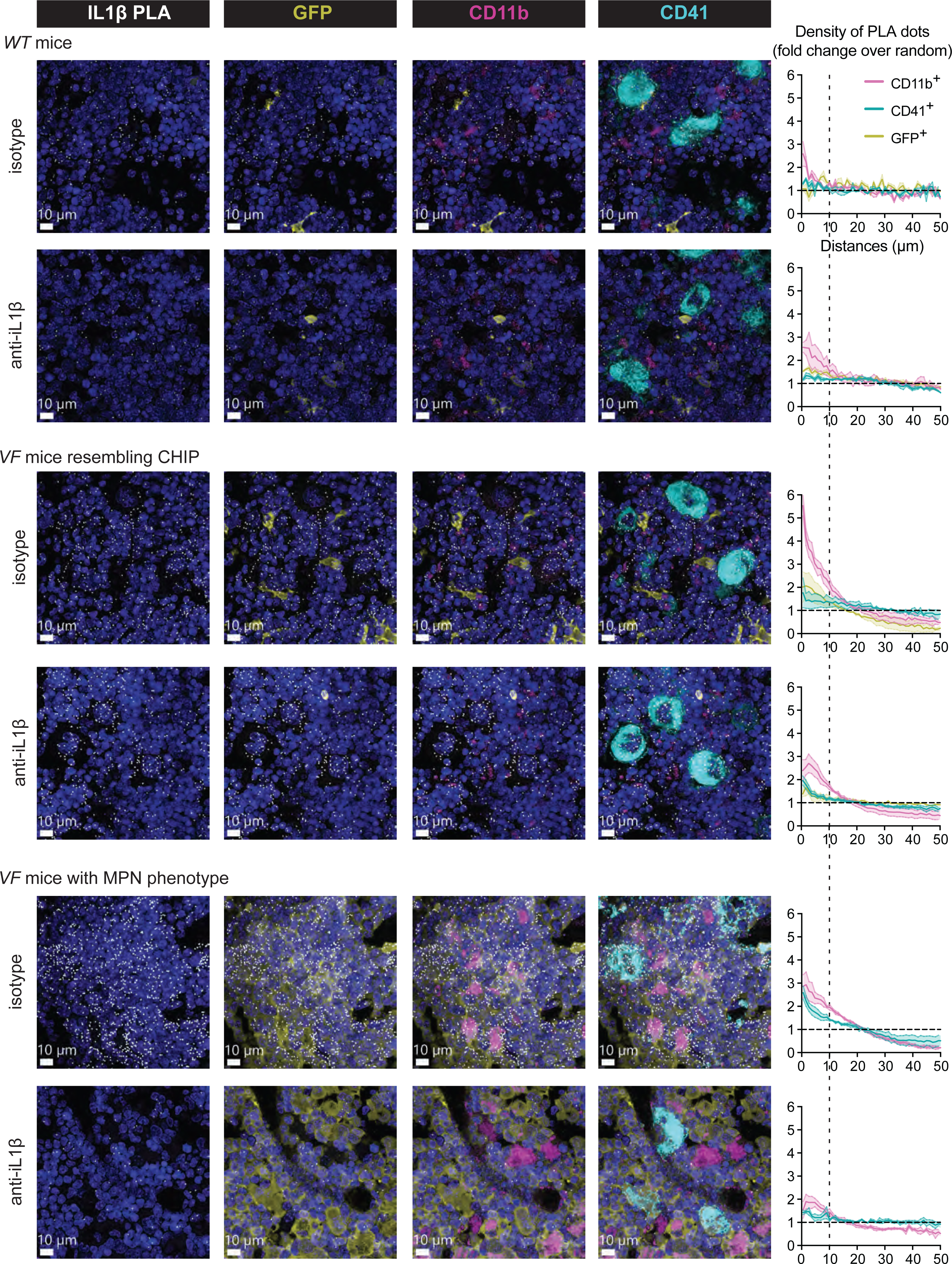
In situ bone marrow 3D fluorescence imaging with Proximity Ligation Assay (PLA) for IL-1β single molecule visualization. Representative images show the distribution of the IL-1β PLA signals relative to CD11b+ myeloid cells and CD41+ megakaryocytes in BM sections of *VF* and *WT* mice (Figure 6). The 3D spatial densities of PLA signals relative to the distances of the closest CD11b+ myeloid cells, CD41+ megakaryocytes or GFP-positive *JAK2*-mutant cells are shown in the graphs next to the pictures.

In *VF* mice, the IL-1β PLA signals were also enriched in the vicinity of CD41+ megakaryocytes and these signals were strongly reduced in mice treated with anti-IL-1β antibody (Figure 6). These results show that *JAK2*-V617F upregulates IL-1β expression not only in myeloid cells, but also in megakaryocytes and suggest that the treatment with anti-IL-1β antibody interrupts the feedback activation of IL-1β. Consistent with the IL-1β PLA data, expression of *IL-1β* mRNA was also increased in sorted monocytic and megakaryocytic cells from BM of *VF* mice compared to WT mice (Supplemental Figure S9B and S9C).

### *IL1B* polymorphisms are associated with increased serum IL-1β levels in MPN patients

Since IL-1β serum levels were increased in patients with MPN and correlated with the *JAK2*-V617F VAF (22), we examined whether genetic polymorphisms located in the *IL1B* gene correlated with higher IL-1β serum levels in MPN patients (Figure 7). Several functionally relevant polymorphisms in *IL1B* gene have been reported (32, 33). We selected two *IL1B* gene polymorphisms that were associated with increased production of IL-1β in non-hematological cancers (34-36), to compare their frequencies in MPN patients and normal controls (Figure 7A). We found that the frequencies of the homozygous AA genotype of rs16944 (G511>A) located in the promoter region of *IL1B* and of the homozygous GG genotype of rs1143627 (A31>G) located closer to the *IL1B* coding regions were almost 3-fold higher in patients with *JAK2*-V617F positive MPN compared to normal controls. Furthermore, MPN patients with AA and GG genotypes displayed higher IL-1β serum levels than MPN patients with GG/GA or AA/AG genotypes (Figure 7). Although, validation in a larger cohort of MPN patients is needed, our data support the model that presence of IL-1β favors the expansion of the *JAK2*-mutant clone.

**Figure 7.**
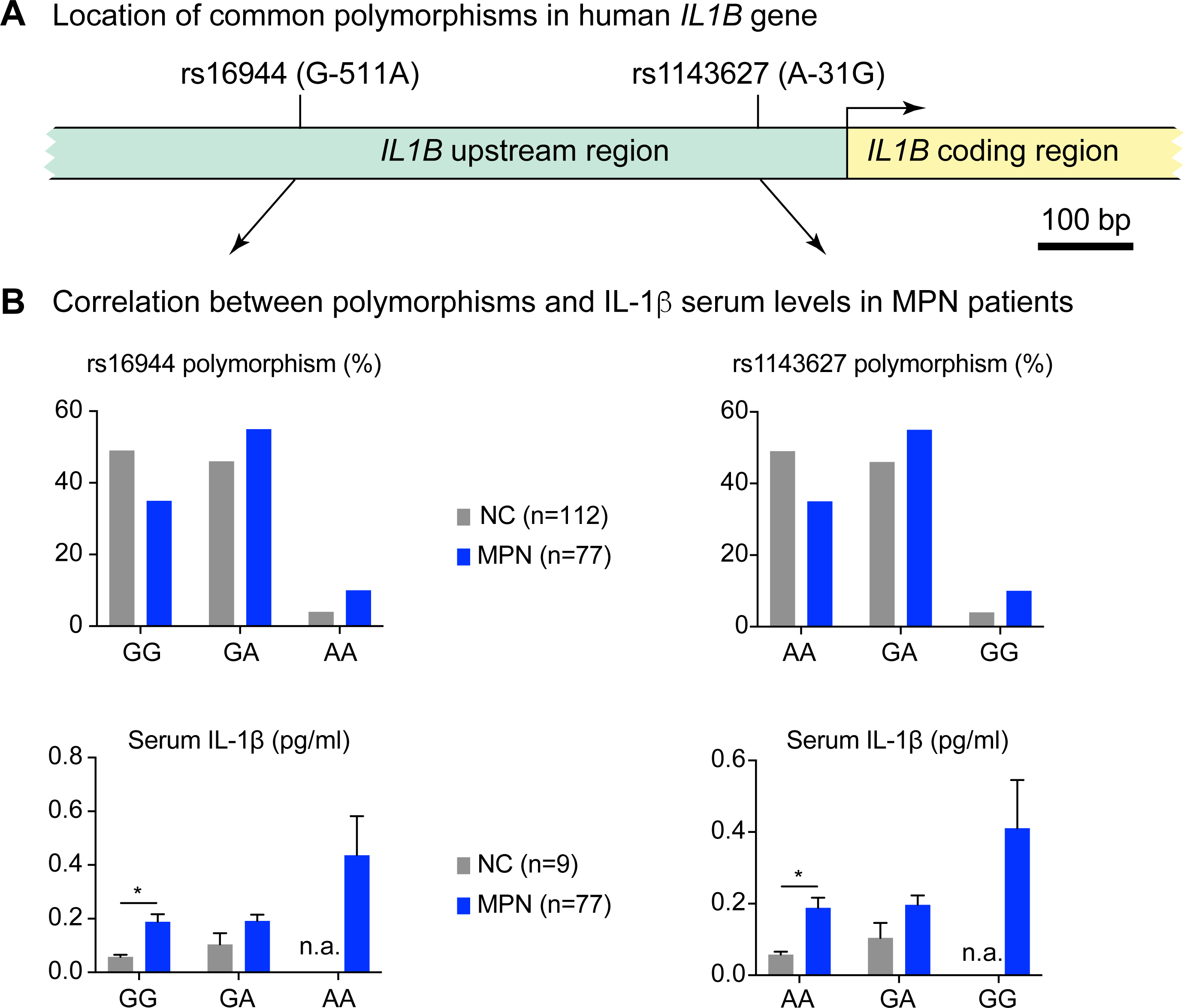
*IL1B* polymorphisms are associated with increased serum IL-1β levels in MPN patients. **(A)** Location of two most commonly studied *<u>IL1B</u>* SNPs in the *IL1B* gene. **(B)** Bar graph showing the frequency of genotypes for *IL1B* SNP rs16944 (-511G/A) and serum IL-1β levels of individuals with different genotypes in normal controls (NC) and MPN patients (left panel). Bar graph showing the frequency of genotypes for *IL1B* SNP rs1143627 (-511G/A) and serum IL-1β levels of individuals with different genotypes in normal controls (NC) and MPN patients (right panel). Two-tailed unpaired t tests were performed for statistical comparisons. All data are presented as mean ± SEM. *P < .05; **P < .01; ***P < .001; ****P < .0001.

## Discussion

In this study, we investigated whether inflammation mediated by IL-1β promotes the early expansion of the *JAK2*-V617F mutant clone and can thereby favor the conversion from a CHIP-like state to MPN. Competitive BM transplantations at high dilutions allowed us to examine oligo- or monoclonal MPN disease initiation starting from a small number (1-3) LT-HSCs per transplanted mouse (37). Genetic ablation of *IL-1β* from *JAK2*-mutant donor HSCs significantly reduced the frequency of engraftment and also lowered the frequency of conversion from CHIP to MPN among mice that engrafted (Figure 1). HSCs expressing *VF*, but deficient for *IL-1β* were also less functional in secondary transplantations, consistent with a role of IL-1β for optimal HSC expansion and long-term repopulation capacity (Figure 2). Although the complete genetic loss of *IL-1β* in donor and recipients eliminated the differences in engraftment and MPN disease initiation compared to *VF* (Supplemental Figure S2), treatment with anti-IL-1β antibody decreased engraftment and MPN disease initiation in competitive BM transplantations at high dilution (Figure 4). Our data establish *IL-1β* as an important trigger for the conversion from a CHIP-like state to MPN in carriers of *JAK2*-V617F.

Our data obtained with the *IL-1R1* knockout are in line with the results of *IL-1β* knockout. Since the *IL-1R1* is required for the positive autocrine feedback loop that augments IL-1β production (18, 19, 38), loss of *IL-1R1* in *VF*;*IL-1R1^-/-^* hematopoietic cells prevented the massive overproduction of IL-1β observed in *VF* hematopoietic cells. IL-1β produced by the *VF*;*IL-1R1^-/-^* hematopoietic cells was still adequate to assure high percentage of engraftment, but the levels were not sufficient to reach the same high frequency of conversion from CHIP to MPN observed with *VF* hematopoietic cells (Figure 3A). Loss of *IL-1R1* in the recipients reduced both the frequency of engraftment and the conversion rate from CHIP to MPN, suggesting that alterations in the non-hematopoietic microenvironment explain the effects of IL-1β on engraftment and MPN disease initiation. Indeed, we found that *IL-1R1* expression in non-hematopoietic BM stromal cells was required for the damaging effects of IL-1β produced by *JAK2*-mutant cells (Figure 3C). Consistent with these findings, anti-IL-1β antibodies prevented the reduction of BM MSCs (Figure 5), including HSC niche-forming nestin+ MSCs that are susceptible to IL-1β-induced damage (21, 22). Interestingly, the reduction of BM MSCs was also observed in mice engrafted with *VF* BM that developed a CHIP-like state, but did not progress to MPN (Figure 5), suggesting that the microenvironmental niche may regulate the size of the *JAK2*-mutant HSC pool. The mechanism of how the presence of nestin+ MSCs interferes with the expansion of the *JAK2*-V617F clone is currently unknown and will be the subject of future studies. We found that in addition to myeloid cells, previously shown to be the main IL-1β producing cells (39), megakaryocytes also contribute to the IL-1β overproduction in *VF* mice (Figure 6), as well as to promoting myelofibrosis (22).

Recent studies have reported associations between inflammatory cytokines and the expansion of CHIP clones carrying mutations other than *JAK2*-V617F. TNF-α promoted the expansion of hematopoietic cells with mutations in *Dnmt3a* or *Asxl1* (40, 41), and IL-1α and IL-1β mediated expansion of hematopoietic clones carrying *Tet2* mutations (42). As such, inhibiting inflammation could decrease the probability of MPN disease initiation from *JAK2*-mutant HSCs. Although aspirin slightly reduced the frequency of engraftment of *VF* BM, as a monotherapy it had no effect on the frequency of MPN initiation (Supplemental Figure S7). Aspirin in combination with anti-PD1 showed therapeutic benefit in a *Braf-V600E* melanoma model (43). *JAK2*-V617F clones induce a strong IL-1-mediated response, as seen by the relatively high levels of IL-1β and IL-6 in plasma as compared to other cytokines such as IFN-γ or KC/GRO. IL-1β antibody neutralized IL-1β and substantially reduced MPN initiation. The anti-IL-1β antibody canakinumab is currently being tested in a clinical trial for clonal cytopenias of unknown significance (CCUS) (https://clinicaltrials.gov/ct2/show/NCT05641831).

Our results provide evidence that *JAK2*-mutant clone derived IL-1β production generates an inflammatory environment in the BM that promotes *JAK2*-V617F clonal expansion and initiation of MPN disease. Inhibition of IL-1β using anti-IL-1β antibodies may be beneficial in individuals with *JAK2*-V617F CHIP who are at increased risk of progression to MPN.

## Methods

### Patients and healthy controls

Blood samples and clinical data of MPN patients were collected at the University Hospital Basel, Switzerland. Blood samples from healthy controls were obtained from the local blood donation center (Stiftung Blutspendezentrum SRK beider Basel). The study was approved by the local Ethics Committees (Ethik Kommission Beider Basel). Written informed consent was obtained from all patients in accordance with the Declaration of Helsinki. The diagnosis of MPN was established according to the World Health Organization and International Consensus Classification criteria (44, 45). Information on diagnosis and gene mutations of MPN patients included in this study are specified in Supplemental Table 1.

### Mice

We used conditional *JAK2*-V617F transgenic mice (46), *SclCre^ER^* mice (47), *UBC-GFP* transgenic mice (48). *IL-1β* knockout mice (49), and *IL-1R1* knockout mice (50). Tamoxifen inducible *SclCre^ER^;V617F (VF)* mice were characterized previously (51). Cre^ER^ in these mice was activated by intraperitoneal injection of 2 mg tamoxifen (Sigma Aldrich) for 5 consecutive days. All mice were of pure C57BL/6N background, and kept under specific pathogen-free conditions with free access to food and water in accordance with Swiss federal regulations. All animal experiments were approved by the Cantonal Veterinary Office of Basel-Stadt, Switzerland.

### Bone marrow transplantations and drug treatment

BM cells from transgenic mice were harvested 6-8 weeks after tamoxifen induction, mixed with wildtype BM competitor cells and injected into the tail vein of lethally irradiated recipient mice (12 Gy). Blood was drawn from tail vein every 4–6 weeks. GFP-chimerism was assessed by flow cytometry and complete blood counts (CBC) were performed on Advia120 Hematology Analyzer with Multispecies Version 5.9.0-MS software (Bayer). Aspirin (Sigma) was given in the drinking water (150 µg/ml) and water with aspirin was changed every 3-4 days. Mouse IgG2a anti-mouse IL-1β antibody (01BSUR; Novartis Pharma AG, Basel, Switzerland) (52-54), or mouse IgG2a isotype were injected intraperitoneally once per week (10 mg/kg*qw, i.p.).

### Flow cytometry and fluorescence activated cell sorting

BM cells were harvested from long bones (2 tibias and 2 femurs) by crushing bones in staining media (Dulbecco’s PBS+ 3% FCS+ pen/strep). Cells were filtered through 70μm nylon mesh to obtain a single-cell suspension. Total spleen cells were harvested by crushing the spleen against 100 μm cell strainer. Red blood cells were lysed (ACK buffer, Invitrogen) and stained with following antibodies for FACS analysis: A mixture of biotinylated monoclonal antibodies CD4, CD8, B220, TER-119, CD11b, and Gr-1 was used as the lineage mix (Lin) together with Sca-1-APC-Cy7, c-kit-BV711, CD48-AF700, CD150-PE-Cy7, CD16-PE, CD41-BV605, CD105-PerCP-Cy5.5 all from BioLegend and CD34-AF647 (BD biosciences) were used as primary antibodies. Cells were washed and stained with streptavidin pacific blue secondary antibody (Invitrogen). Mouse stromal cells were obtained by crushing mouse bones directly in 0.25% collagenase I in 20% FBS/PBS solution and digesting bones and cells at 37°C water bath for 45 minutes. Cells were filtered through 70μm nylon mesh, red blood cells were lysed (ACK buffer, Invitrogen) and cells were stained with CD45-PE-Cy7, CD31-PerCP-Cy5.5, TER-119-APC, Sca-1-APC-Cy7, PDGFRα-PE from BioLegend. Sytox-Blue or Green (Invitrogen) was used to exclude dead cells during FACS analysis. Live, singlet cells were selected for gating. Cells were analyzed on a Fortessa Flow Cytometer (BD biosciences). Data were analyzed using FlowJo (version 10.7.1) software. Mouse BM populations including monocytes, granulocytes and megakaryocytes were FACS sorted directly in PicoPure RNA extraction buffer (Applied Biosystems). Sorting and gating strategy is described in Supplemental Figure S8.

### RNA isolation and qPCR

RNA from FACS sorted monocytes, megakaryocytes and granulocytes from bone marrow of *WT* and *VF* mice were prepared using PicoPure RNA isolation kit (Applied Biosystems). RNA was then reverse transcribed to cDNA using High-Capacity cDNA Reverse Transcription Kit from Applied Biosystems according to manufacturer’s instructions. *IL-1β* and *Gapdh* gene expression in mouse bone marrow populations were quantified by TaqMan gene expression assay (Assay ID: and; ThermoFisher Scientific). Each sample was run in triplicates using 2-5 ng cDNA in a 384 well plate and the qPCR was performed using VIIA 7 real time PCR instrument from Applied Biosystems.

### Cytokine analysis

Mouse blood was drawn by cardiac puncture and collected in ETDA tubes. Platelet depleted plasma was collected by centrifuging the blood at 5000g for 20 minutes at 4°C. Mouse BM lavage was prepared by flushing one femur and one tibia with 500µl ice-cold PBS and centrifuging the cell suspension at 300g for 10 minutes at 4°C. IL-1β and other pro-inflammatory cytokine levels in mouse BM and plasma were measured by ELISA kits from R&D systems and Mesoscale Discovery according to manufacturer’s instructions. Single analyte data was plotted in GraphPad Prism software using an XY data table and the standard curve was analyzed using a sigmoidal 4-PL equation and the values of unknowns were interpolated. Multiplex cytokine data from a 96-well plate was read using Mesoscale Meso Sector S 600 instrument and the data was analyzed with Discovery Workbench 4.0 software.

### Histology

Bones (sternum and/or femur), spleens and livers were fixed in paraformaldehyde, embedded in paraffin and tissue sections were stained with H&E, or Gömöri. Pictures were taken with 10x, 20x and 40x objective lens using Nikon Ti inverted microscope and NIS Software. All histological assessments of BM, spleen and liver were performed by an experienced hemato-pathologist who had no information about the genotypes or the treatment modalities. For staining of nerve fibers and Schwann cells, mouse skull bones were fixed in 2% formaldehyde/PBS solution for 2 hours at 4°C, washed with PBS and stored in PBS at 4°C until further analysis. Femurs were fixed on a shaker with 2% formaldehyde/PBS solution for 24 hours at 4°C, then washed with PBS and decalcified with 250mM EDTA solution for 10-12 days at 4°C. Decalcified bones were transferred to 30% sucrose/PBS solution for 24 hours and then to 50% OCT and 50% (30% sucrose/PBS) solution for another 24 hours. Bones were then embedded in OCT and kept at −80°C until cryostat sections and the whole mount staining of the skull bones were performed. The antibodies used for immunofluorescence staining were anti-TH (Rabbit pAb, Millipore) and anti- GFAP (Rabbit pAb, Dako). Confocal images were acquired with a laser scanning confocal microscope (Zeiss LSM 700). At least 3 different sections were used for quantification using ImageJ software.

### Immunofluorescence and Proximity Ligation Assay

Proximity Ligation Assay (PLA) and immunofluorescence staining of femur sections were performed as previously described.(30) Briefly, femurs were fixed in 4% methanol-free formaldehyde (Thermo Scientific) for 24 hours at 4 °C post dissection. After 14-day decalcification in 10% EDTA (BioSlove, pH 8), femurs were embedded in 4% low-gelling temperature agarose (Merck) and sectioned with a vibratome (Leica VT1200 S) into 150 μm thick sections. Staining steps were performed at room temperature (20-25°C) on superfrost glass slides (VWR) in double adhesive GeneFrame Chambers (Thermo Scientific) with gentle rocking. Sections were blocked and permeabilized with 10% donkey serum (Jackson ImmunoResearch), 1% Triton X-100 (PanReac AppliChem), 0.05% Tween-20 (Merck), 20% DMSO (Merck) in TBS (0.1M Tris, 0.15M NaCl, pH 7.5) for 2 hours minimum. Primary PLA antibodies, chicken polyclonal anti-GFP (Aves; GFP-1020), rabbit polyclonal anti-IL1-β (Invitrogen; P420B), rat IgG2b monoclonal anti-CD11b (Invitrogen; 14-0112-82), rat IgG1 monoclonal Alexa Fluor 647 anti-CD41(Biolegend; 133934) and secondary antibodies, donkey polyclonal anti-chicken IgY AF488 (Invitrogen; A78948), donkey polyclonal anti-rabbit IgG (H+L) (Merck; DUO92005-100RXN), donkey polyclonal anti-rabbit IgG (H+L) (Merck; DUO92002-100RXN), mouse monoclonal anti-rat IgG2b (Biolegend; 408207) were applied for 2 hours. In addition, 4 washing steps, 30 minutes each, were performed between incubations. Duolink *In Situ* detection reagents Orange (Merck; DUO2007-100RXN) was used for IL1-β single molecule visualization.

### Confocal Microscopy, Image Visualization and Analyses

Images were acquired on a Leica TCS SP8 confocal microscopy using Leica type G immersion liquid with a 63x glycerol immersion lens (NA 1.3, FWD 0.28 mm). The scanning was performed at 400 Hz at room temperature in bidirectional mode at 1024×1024 pixel resolution. Only HyD detectors were used for signal acquisition. Images shown were the MIP (maximum intensity projection) from Imaris 3D reconstruction. Isosurface (CD41+ megakaryocytes and CD11b+ monocytes) and spot (IL-1β PLA) were generated, and 3D distance-transformed in Imaris v9.9.1. PLA signals in the bone, blood vessels, and nucleus were masked out. Random dots were sampled 100 times with the same number as IL-1β PLA dots, resulting in an average dot count every 1μm increment from the Isosurface. The enrichment profiles towards segmented structures were created from the ratio of IL-1β PLA dot to random dot count per distance bin.

## Statistical analyses

The unpaired two-tailed Student’s t-test analysis was used to compare the mean of two groups. Normality tests were performed to test whether the data follows a normal distribution. When the distribution was not normal, non-parametric Mann-Whitney t-tests were performed. For samples with large variances, Welch’s correction was applied for t-test. Multiple t-tests with or without correction or one-way or two-way ANOVA analyses followed by Dunn’s or Tukey’s comparison tests for multiple group comparisons followed by Bonferroni’s correction of p-values. Data were analyzed and plotted using Prism software version 7.0 (GraphPad Inc.). All data are represented as mean ± SEM. Significance is denoted with asterisks (*p<0.05, **p<0.01, ***p < 0.001, ****p < 0.0001).

## Data availability

For original data and reagents, please contact.

## Author contributions

SR designed and performed the research, analyzed data, and wrote the manuscript; YZ and EG performed research and analyzed data, NH, QK, CS, MU and HHS performed research; MSB and JRP analyzed clinical data; SD performed and analyzed histopathology of mouse tissues; CF, TS and SMF designed research and analyzed data; RCS designed the research, analyzed the data, and wrote the manuscript.

## Acknowledgments

The authors thank the members of the Flow cytometry core facility of the DBM for FACS experiments, and laboratory members for helpful discussions and critical reading of the manuscript. This work was supported by grants from the Swiss National Science Foundation (31003A_166613, and 310030_185297) and Swiss Cancer Research (KFS-3655-02-2015 and KFS-4462-02-2018) to RCS, and core support grants from MRC to the Cambridge Stem Cell Institute; National Health Service Blood and Transplant (United Kingdom), European Union’s Horizon 2020 research (ERC-2014-CoG-648765), MRC-AMED grant MR/V005421/1 and a Programme Foundation Award (C61367/A26670) from Cancer Research UK to SMF.

