## Supplemental Material for "IL-1β promotes MPN disease initiation by favoring early clonal expansion of *JAK2*-mutant hematopoietic stem cells"

**Supplemental Figure S1 (related to Figure 1):** Competitive transplantations at 1:100 dilution into wildtype (*WT*) recipients with *WT* competitor cells

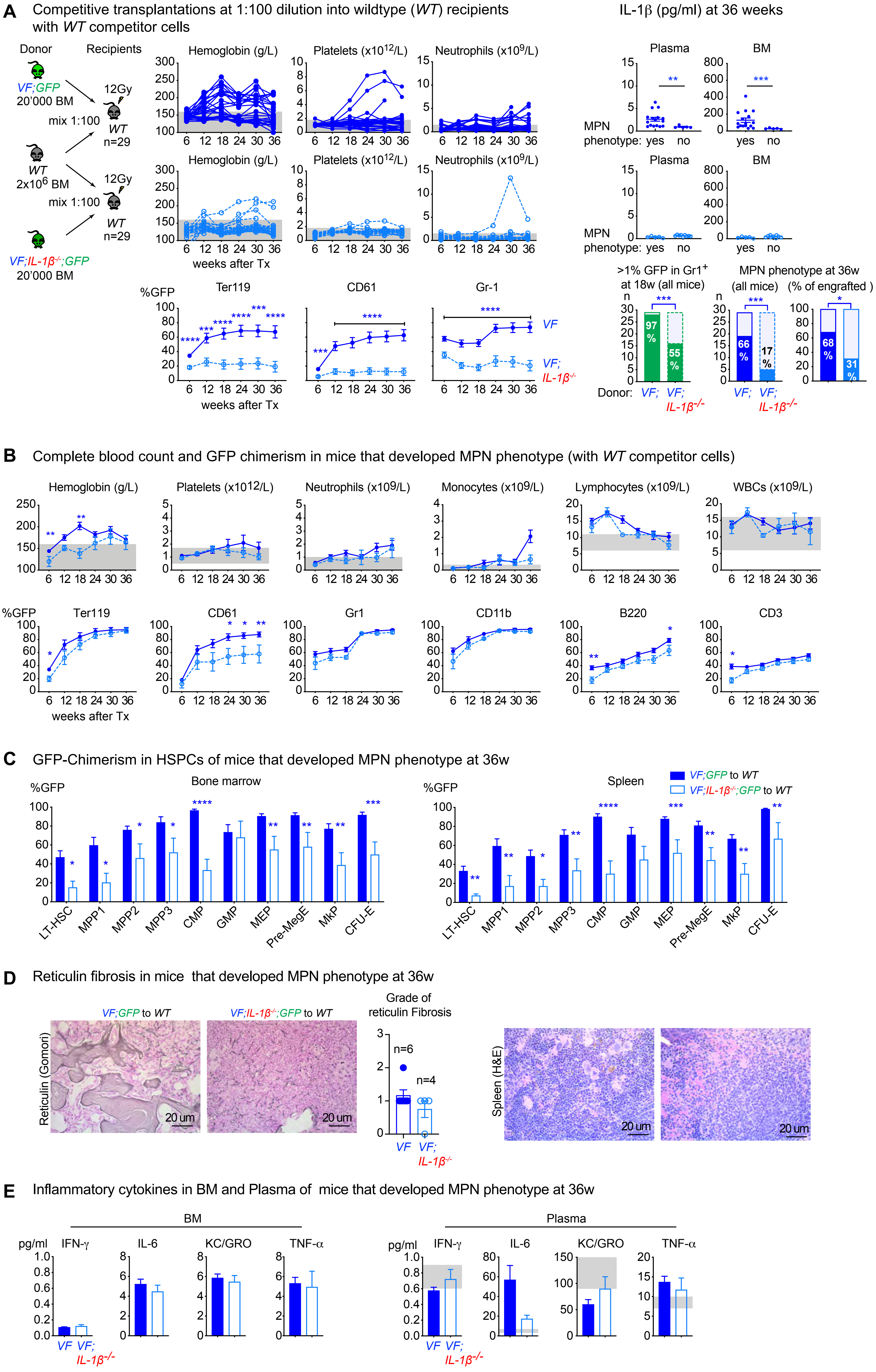

#### Legend to Supplemental Figure S1 (related to Figure 1)

**Supplemental Figure S1. Loss of *IL-1β* from *JAK2*-mutant hematopoietic cells reduces MPN disease initiation (Transplantation at 1:100 dilution into *WT* recipients with BM competitor cells from a *WT* donor). (A)** Schematic drawing of the experimental setup for competitive transplantation at 1:100 dilution. Bone marrow (BM) from *VF;GFP* or *VF;IL-1β-/-;GFP* donor mice was mixed with a 100-fold excess of BM competitor cells from a *WT* donor. The time course of blood counts from individual mice that received BM from *VF;GFP* (upper panel) or *VF;IL-1β-/-;GFP* donors (middle panel), and the GFP chimerism in peripheral blood (lower panel) are shown. Multiple t tests were performed for statistical analyses. IL-1β protein levels in plasma and BM lavage (1 femur and 1 tibia) of mice with or without MPN phenotype is shown (right panel). Non-parametric Mann-Whitney two-tailed t test was performed for statistical comparisons. Bar graphs (bottom right) show the percentages of mice that showed engraftment defined as GFP-chimerism >1% at 18 weeks after transplantation and the percentages of mice that developed MPN phenotype (elevated hemoglobin and/or platelet counts). p values in lower panel were computed using Fisher's exact test. **(B)** Time course of mean blood counts and GFP chimerism in peripheral blood of *WT* mice transplanted with BM from *VF;GFP* or *VF;IL-1β-/-;GFP* and *WT* competitor cells that developed MPN phenotype during 36-weeks follow-up. Multiple t tests were performed for statistical analyses. **(C)** GFP-chimerism in hematopoietic stem and progenitor cells (HSPCs) at 36 weeks after transplantation in BM (left) and spleen (right) of *WT* mice transplanted with BM from *VF;GFP* or *VF;IL-1β-/-;GFP* and *WT* competitor cells that developed MPN phenotype. Multiple t tests were performed for statistical analyses. **(D)** Representative images of reticulin fibrosis staining in BM (left panel) and H&E staining in spleen (right panel) of mice showed MPN phenotype at 36 weeks after transplantation. Histological grade of reticulin fibrosis in BM is shown in a bar graph (right). **(E)** Levels of Inflammatory cytokines in BM lavage (1femur and 1 tibia) and plasma of mice that displayed MPN phenotype at 36 weeks after transplantation. Grey shaded area represents normal range. All data are presented as mean ± SEM. *P < .05; **P < .01; ***P < .001; ****P < .0001.

#### Supplemental Figure S2 (related to Figure 1C)

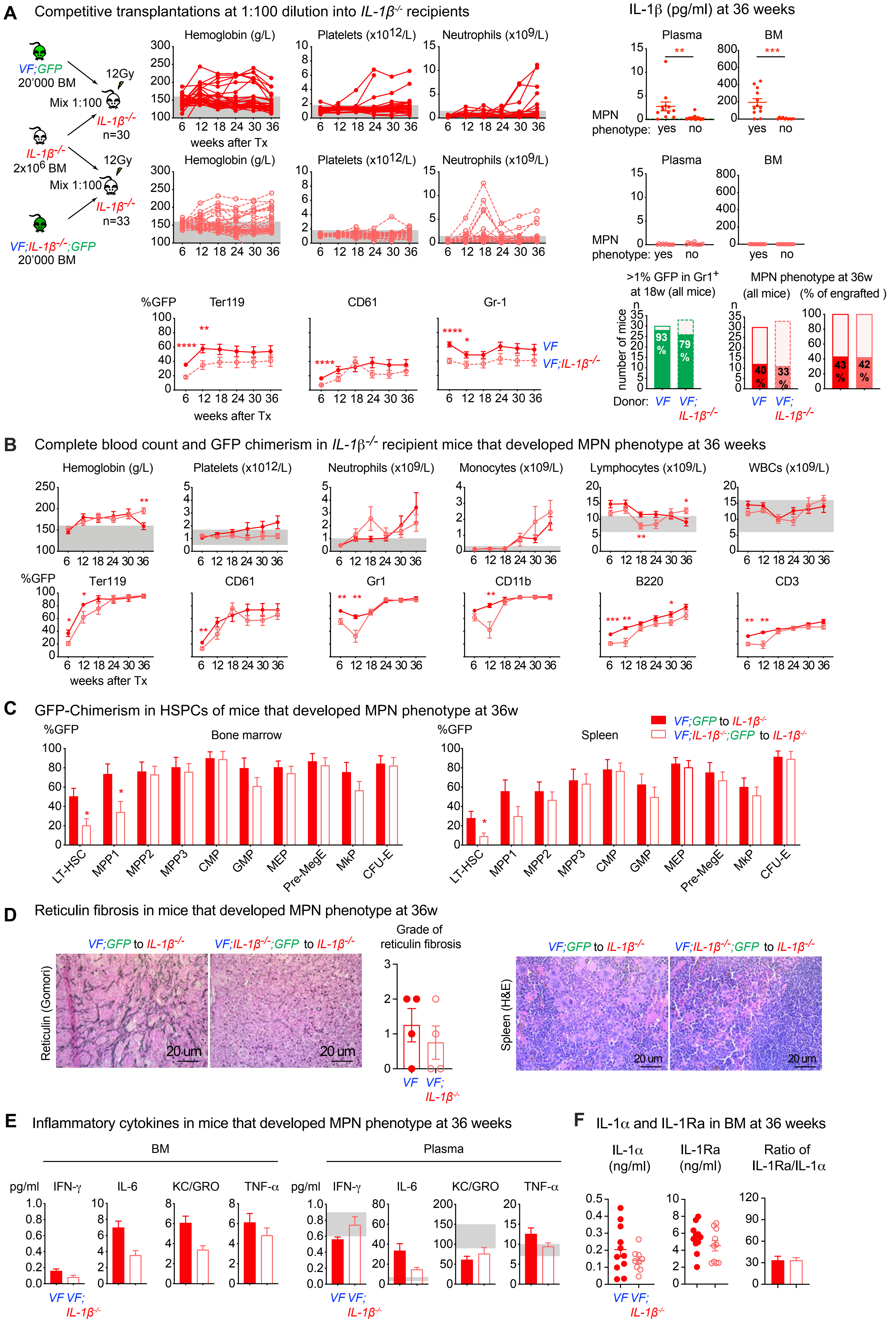

#### Legend to Supplemental Figure S2 (related to Figure 1)

**Supplemental Figure S2. Loss of *IL-1β* from *JAK2*-mutant hematopoietic cells reduces MPN disease initiation (Transplantation at 1:100 dilution into *IL-1β*$^{-/-}$ recipients with *IL-1β*$^{-/-}$ competitor cells). (A)** Schematic drawing of the experimental setup for competitive transplantation at 1:100 dilution into *IL-1β*$^{-/-}$ recipients. Bone marrow (BM) from *VF;GFP* or *VF;IL-1β-/-;GFP* donor mice was mixed with a 100-fold excess of BM competitor cells from *IL-1β-/-* donor. The time course of blood counts from individual mice that received BM from *VF;GFP* (upper panel) or *VF;IL-1β-/-;GFP* donors (middle panel), and the GFP chimerism in peripheral blood (lower panel) are shown. Multiple t tests were performed for statistical analyses. IL-1β protein levels in plasma and BM lavage (1 femur and 1 tibia) of mice with or without MPN phenotype is shown (right panel). Non-parametric Mann-Whitney two-tailed t test was performed for statistical comparisons. Bar graphs (bottom right) show the percentages of mice that showed engraftment defined as GFP-chimerism >1% at 18 weeks after transplantation and the percentages of mice that developed MPN phenotype (elevated hemoglobin and/or platelet counts). p values in lower panel were computed using Fisher's exact test. **(B)** Time course of mean blood counts and GFP chimerism in peripheral blood of *IL-1β*$^{-/-}$ mice transplanted with BM from *VF;GFP* or *VF;IL-1β-/-;GFP* and *IL-1β*$^{-/-}$ competitor cells that developed MPN phenotype during 36-weeks follow-up. Multiple t tests were performed for statistical analyses. **(C)** GFP-chimerism in hematopoietic stem and progenitor cells (HSPCs) at 36 weeks after transplantation in BM (left) and spleen (right) of *IL-1β*$^{-/-}$ mice transplanted with BM from *VF;GFP* or *VF;IL-1β-/-;GFP* and *IL-1β*$^{-/-}$ competitor cells that developed MPN phenotype. Multiple t tests were performed for statistical analyses. **(D)** Representative images of reticulin fibrosis staining in BM (left panel) and H&E staining in spleen (right panel) of mice showed MPN phenotype at 36 weeks after transplantation. Histological grade of reticulin fibrosis in BM is shown in a bar graph (right). **(E)** Levels of Inflammatory cytokines in BM lavage (1femur and 1 tibia) and plasma of mice that displayed MPN phenotype at 36 weeks after transplantation. **(F)** IL-1α and IL-1Ra levels (pg/ml) in BM of mice (from Supplemental Figure S2A) that showed MPN phenotype. Bar graph showing ratio of IL-1Ra to IL-1α in BM. p value was computed using unpaired two-tailed t-tests with Welch's correction. Grey shaded area represents normal range. All data are presented as mean ± SEM. *P < .05; **P < .01; ***P < .001; ****P < .0001.

#### Supplemental Figure S3 (related to Figure 2)

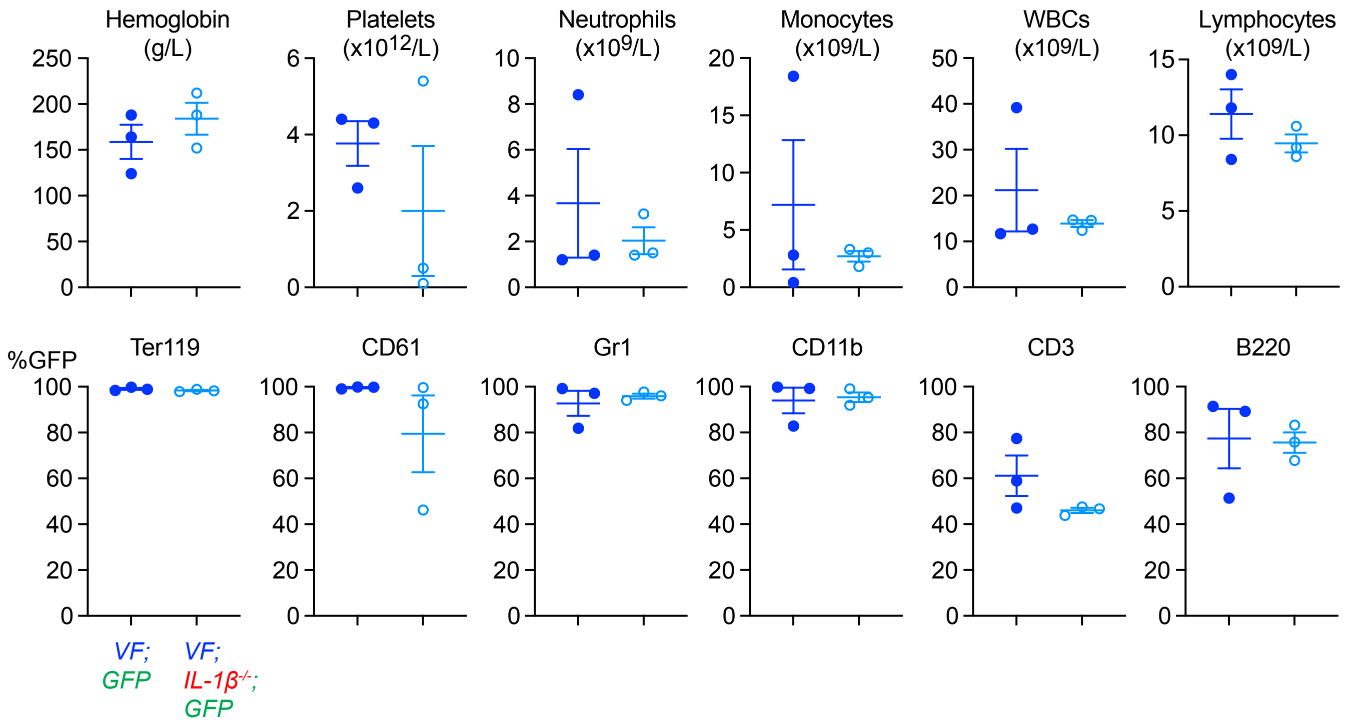


**Supplemental Figure S3.** Blood counts and GFP chimerism of donor mice used for secondary transplantations. Analysis was performed 36 weeks after the initial transplantation. All data are presented as mean ± SEM.

#### Supplemental Figure S4 (related to Figure 2)

**A** Non-Competitive (1:0) secondary transplantations

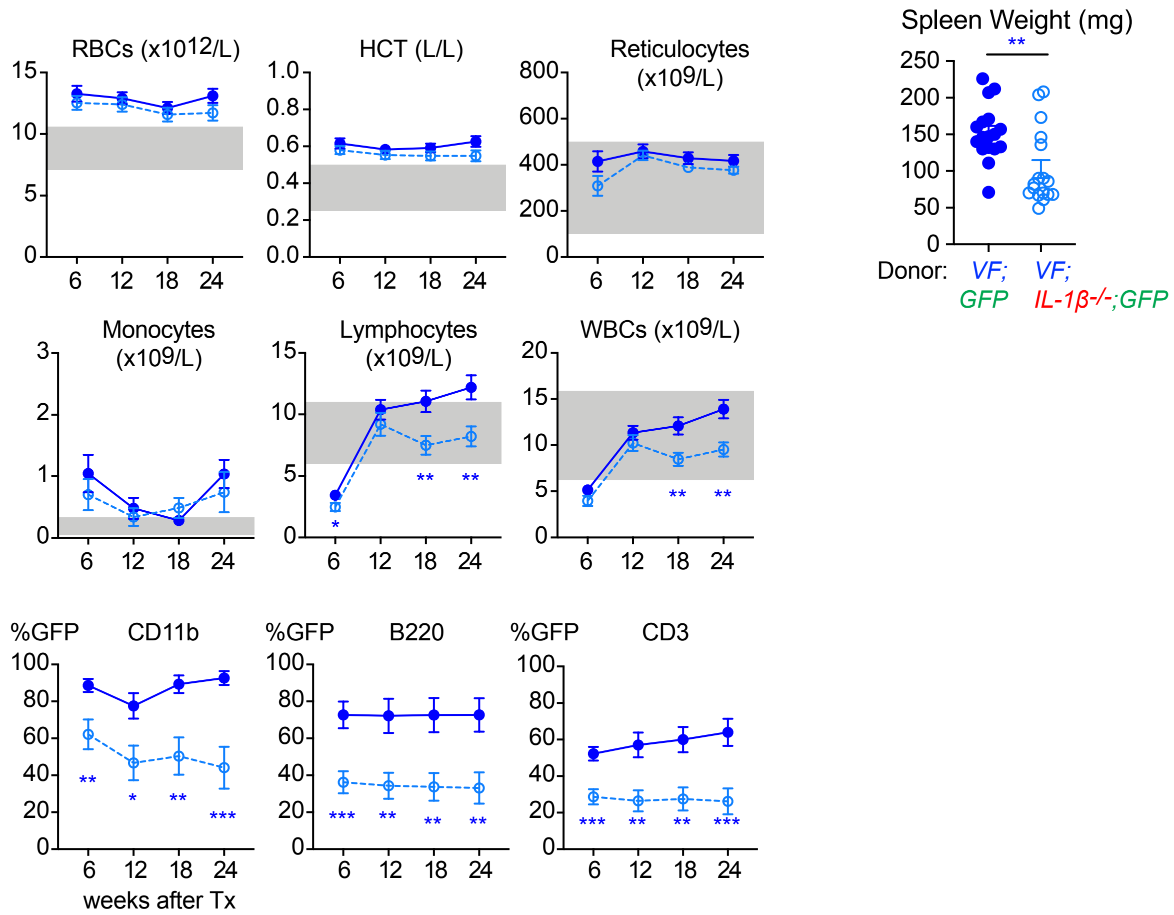


**B** Competitive (1:1) secondary transplantations

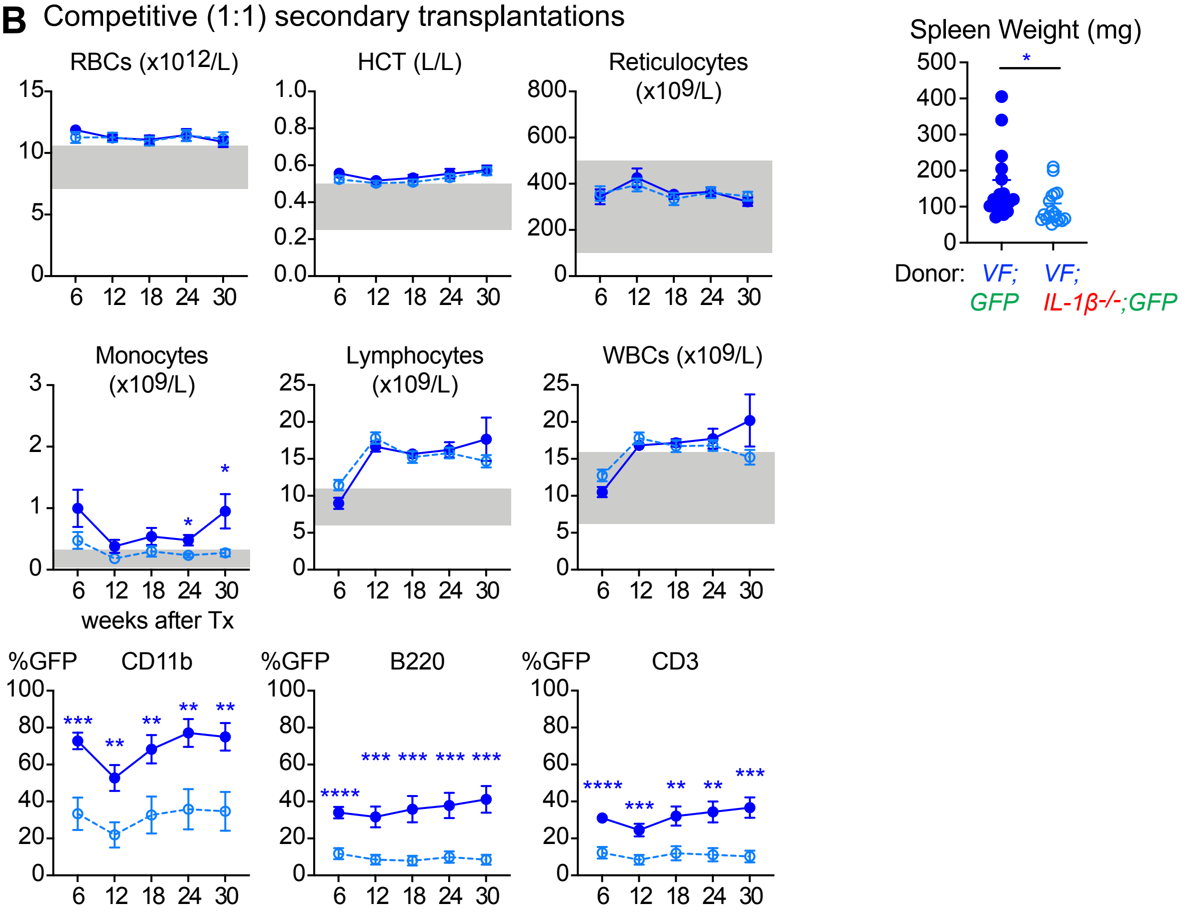


**Supplemental Figure S4. (A)** Red cell parameters and leukocyte counts in secondary transplanted (1:0) mice. Mean GFP-chimerism in CD11b+ monocytes, B220+ B cells and CD3+ T cell lineage is shown. Spleen weight of mice at terminal analysis is shown for both groups. **(B)** Red cell parameters and leukocyte counts in secondary transplanted (1:1) mice. Mean GFP-chimerism in CD11b+ monocytes, B220+ B cells and CD3+ T cell lineage is shown. Spleen weight of mice at terminal analysis is shown for both groups. Grey shaded area represents normal range. All data are presented as mean ± SEM. *P < .05; **P < .01; ***P < .001; ****P < .0001.

**Supplemental Figure S5 (related to Figure 3A):** Competitive transplantations at 1:100 dilution into wildtype (*WT*) recipients

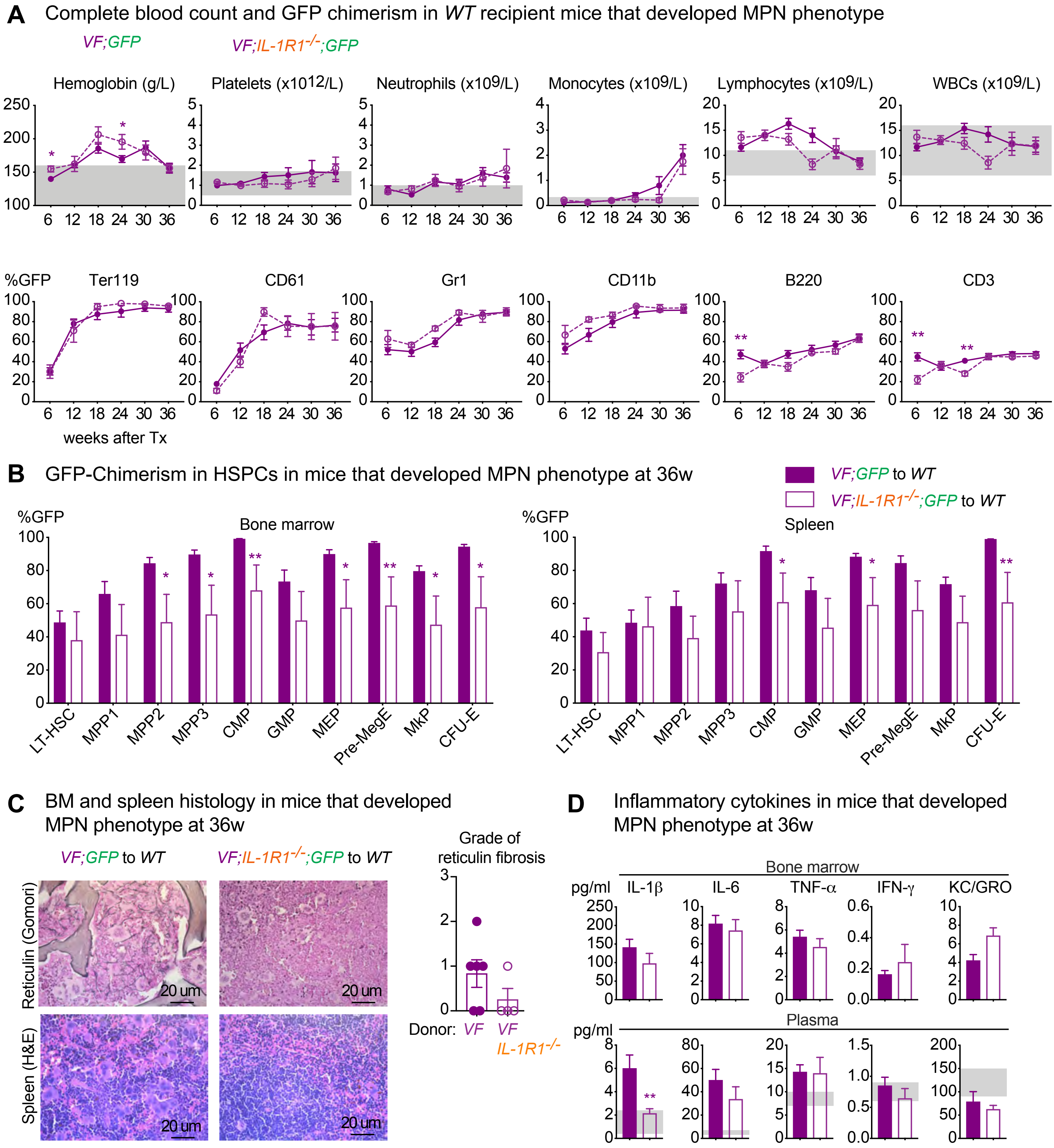


**Supplemental Figure S5. Transplantation at 1:100 dilution into wildtype (WT) recipients with IL-1R1$^{-/-}$ competitor cells.** **(A)** Mean blood counts and GFP chimerism in Ter119, CD61, Gr-1, CD11b (monocytes), B220 (B cells) and CD3 (T cells) in the peripheral blood of WT mice transplanted with BM from VF;GFP or VF;IL-1R1$^{-/-}$;GFP and IL-1R1$^{-/-}$ competitor cells that developed MPN phenotype during 36-weeks follow-up. Multiple t tests were performed for statistical analyses. **(B)** GFP-chimerism in HSPCs in BM (left) and spleen (right) of WT mice transplanted with BM from VF;GFP or VF;IL-1R1$^{-/-}$;GFP and IL-1R1$^{-/-}$ competitor cells that developed MPN phenotype at 36 weeks after transplantation. Multiple t tests were performed for statistical analyses. **(C)** Representative images of reticulin fibrosis staining in BM (upper panel) and H&E staining in spleen (lower panel) of mice that developed MPN at 36 weeks after transplantation. Histological grade of reticulin fibrosis in BM is shown in a bar graph (right). **(D)** Levels of Inflammatory cytokines in BM lavage (1femur and 1 tibia) and plasma of mice that developed MPN at 36 weeks after transplantation. Grey shaded area represents normal range. All data are presented as mean ± SEM. Multiple t tests were performed for statistical analyses. *P < .05; **P < .01; ***P < .001; ****P < .0001.

**Supplemental Figure S6 (related to Figure 3B):** Competitive transplantations at 1:100 dilution into *IL-1R1*$^{-/-}$ recipients

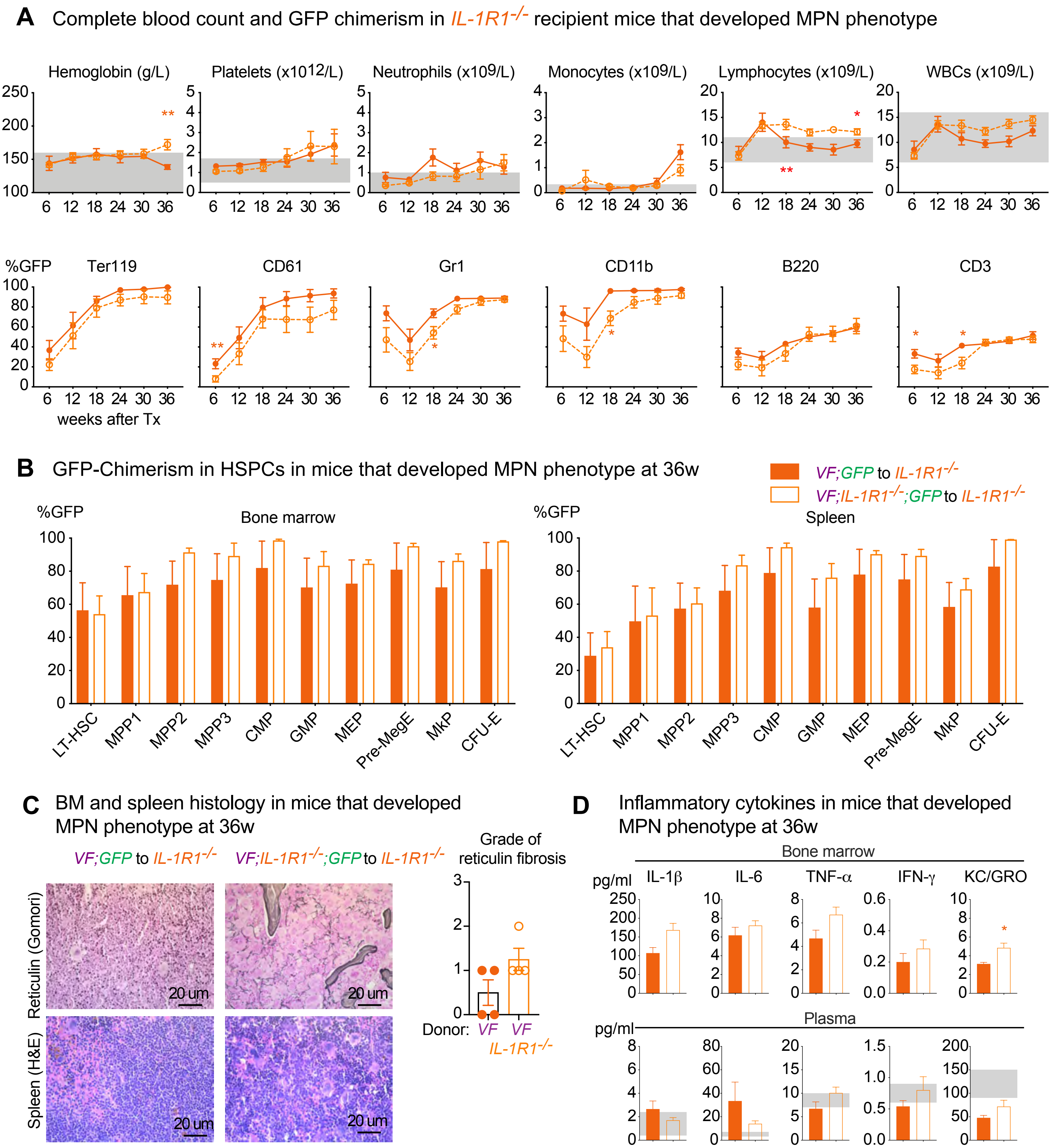


**Supplemental Figure S6. Transplantation at 1:100 dilution into *IL-1R1*$^{-/-}$ recipients with *IL-1R1*$^{-/-}$ competitor cells. (A)** Mean blood counts and GFP chimerism in Ter119, CD61, Gr-1, CD11b (monocytes), B220 (B cells) and CD3 (T cells) in the peripheral blood of IL-1R1$^{-/-}$ mice transplanted with BM from VF;GFP or VF;IL-1R1$^{-/-}$;GFP and IL-1R1$^{-/-}$ competitor cells that developed MPN phenotype during 36-weeks follow-up. Multiple t tests were performed for statistical analyses. **(B)** GFP-chimerism in HSPCs in BM (left) and spleen (right) of IL-1R1$^{-/-}$ mice transplanted with BM from VF;GFP or VF;IL-1R1$^{-/-}$;GFP and IL-1R1$^{-/-}$ competitor cells that developed MPN phenotype at 36 weeks after transplantation. Multiple t tests were performed for statistical analyses. **(C)** Representative images of reticulin fibrosis staining in BM (upper panel) and H&E staining in spleen (lower panel) of mice that developed MPN at 36 weeks after transplantation. Histological grade of reticulin fibrosis in BM is shown in a bar graph (right). **(D)** Levels of Inflammatory cytokines in BM lavage (1femur and 1 tibia) and plasma of mice that developed MPN at 36 weeks after transplantation. Grey shaded area represents normal range. All data are presented as mean ± SEM. Multiple t tests were performed for statistical analyses. *P < .05; **P < .01; ***P < .001; ****P < .0001.

### Supplemental Figure S7

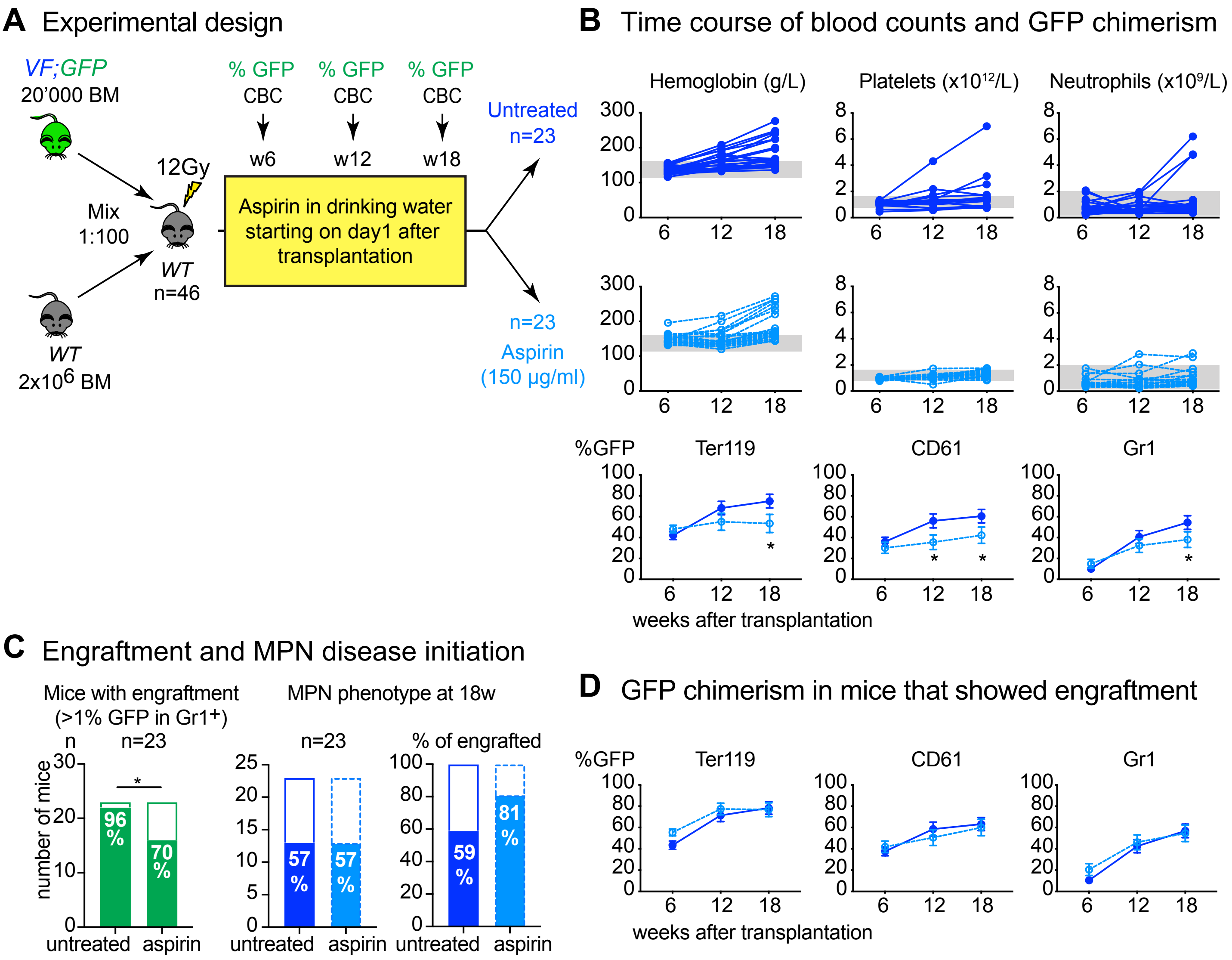


**Supplemental Figure S7. Effects of Aspirin on MPN disease initiation in a mouse model of JAK2-V617F driven clonal hematopoiesis (A)** Schematic drawing of the experimental setup for competitive transplantation at 1:100 dilution. Bone marrow (BM) from a *VF;GFP* donor mouse was mixed with a 100-fold excess of BM competitor cells from a *WT* donor. Mice were randomized into two treatment arms one day after transplantation and treated with either Aspirin or nothing in the driniking water (150 µg/ml) for 18 weeks. **(B)** The time course of blood counts from individual mice for the two treatment arms is shown (top and middle panel). Mean GFP (mutant cell) chimerism in peripheral blood erythroid (Ter119), megakaryocytic (CD61), granulocytic (Gr1) cells are shown in the bottom panel. Multiple t tests were performed for statistical analyses. **(C)** Bar graph showing the percentage of mice that showed engraftment defined as GFP-chimerism >1% at 18 weeks after transplantation. **(D)** Mean GFP (mutant cell) chimerism in peripheral blood of engrafted mice (engraftment defined as GFP-chimerism >1%) at 18 weeks after transplantation. p value in right panel was computed using Fisher's exact test. Grey shaded area represents normal range. All data are presented as mean ± SEM. *P < .05; **P < .01; ***P < .001; ****P < .0001.

#### Supplemental Figure S8 (related to Figure 4)

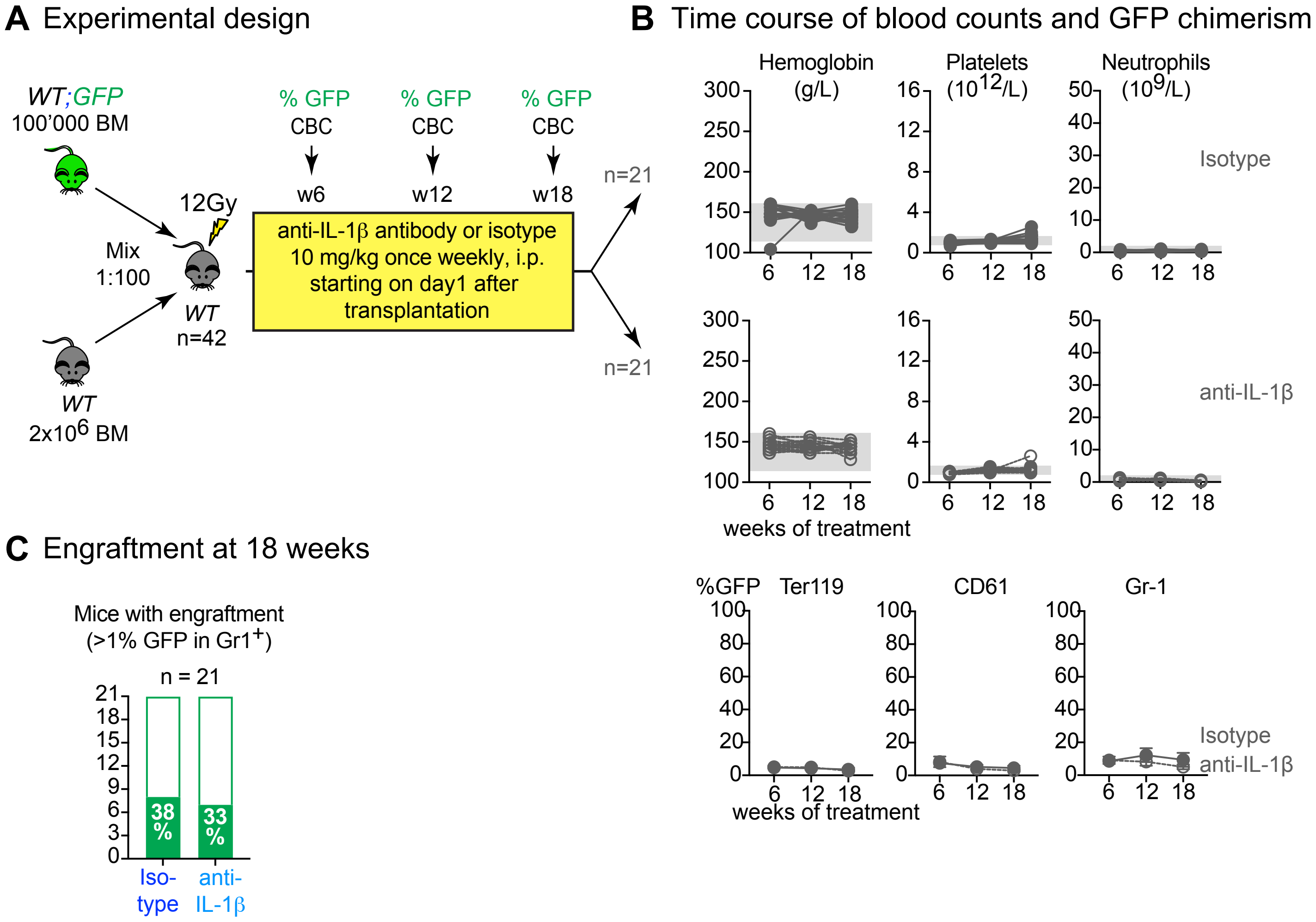


**Supplemental Figure S8. Anti-IL-1β antibody reduces MPN disease initiation in a mouse model of *JAK2*-V617F driven clonal hematopoiesis (A)** Schematic drawing of the experimental setup for competitive transplantation at 1:20 dilution. Bone marrow (BM) from a *WT;GFP* donor mouse was mixed with a 20-fold excess of BM competitor cells from a *WT* donor. Mice were randomized into two treatment arms one day after transplantation and treated with either isotype or anti-IL-1β antibody for 18 weeks. **(B)** The time course of blood counts from individual mice for the two treatment arms is shown (top and middle panel). Mean GFP (mutant cell) chimerism in peripheral blood erythroid (Ter119), megakaryocytic (CD61), granulocytic (Gr1) cells are shown in the bottom panel. Multiple t tests were performed for statistical analyses. **(C)** Bar graph showing the percentage of mice that showed engraftment defined as GFP-chimerism >1% at 18 weeks after transplantation. p value in right panel was computed using Fisher's exact test. Grey shaded area represents normal range. All data are presented as mean ± SEM. *P < .05; **P < .01; ***P < .001; ****P < .0001.

#### Supplemental Figure S9 (related to Figure 6)

**A** Quantifitation of IL-1β PLA dots in bone marrow

**B** Gating strategy for fluorescence activated cell sorting of bone marrow cells

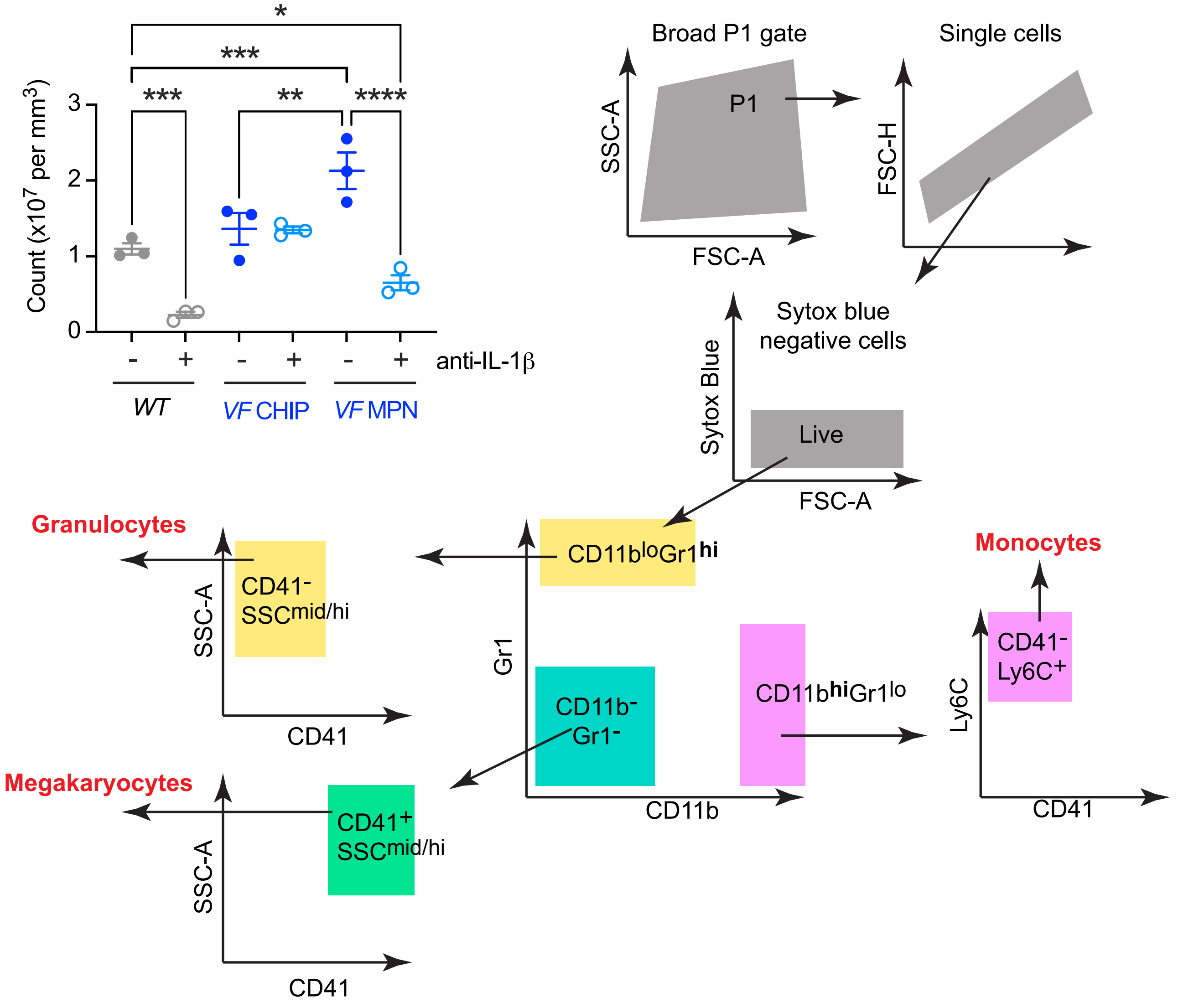


**C** *IL-1β* mRNA expression (TaqMan) in sorted bone marrow cell populations

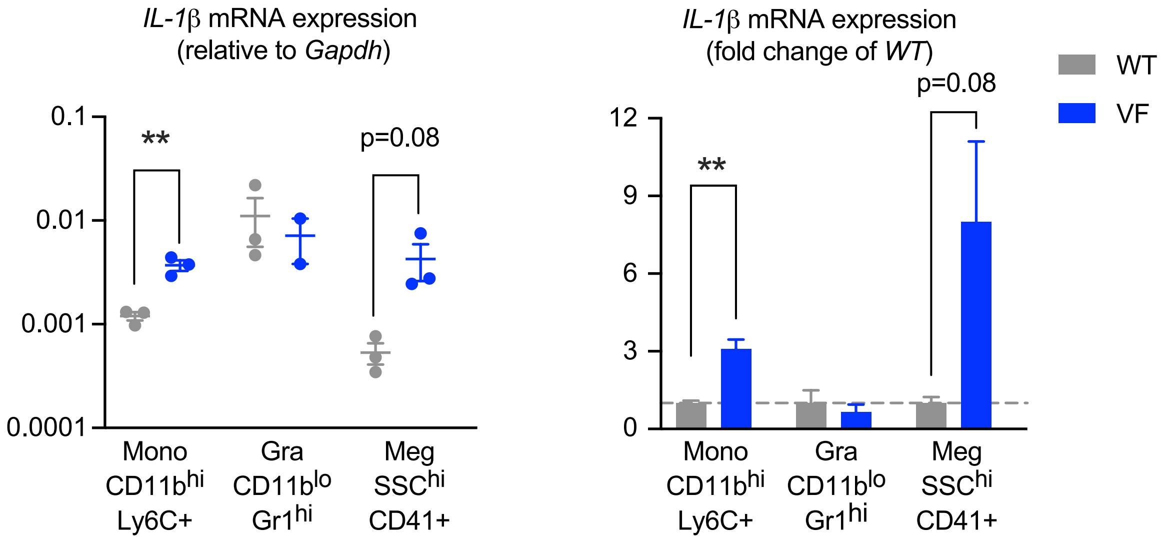


**Supplemental Figure S9**. **Source of IL-1β overproduction in MPN (A)** Quantification of Proximity Ligation Assay (PLA) signals in mouse bone marrow sections. **(B)** Gating strategy for FACS sorting mouse bone marrow populations **(C)** IL-1β mRNA expression in sorted bone marrow cell populations. Two-tailed unpaired t tests were performed for statistical comparisons. All data are presented as mean ± SEM. *P < .05; **P < .01; ***P < .001; ****P < .0001.

**Supplemental Table 1. Characteristics of the MPN patients**

| UPN | Diagnosis | *Jak2*-V617F % VAF | Sex | Additional gene mutations (only if likely pathogenic ) |
|---|---|---|---|---|
| P346A | PV | 79 | female | None |
| P354A | PV | 66 | female | None |
| P355A | PV | 32 | female | None |
| P357A | PV | 43 | male | None |
| P361A | PV | 9 | female | None |
| P362A | PV | 9 | female | None |
| P382A | PV | 41 | male | None |
| P389A | PV | 49 | female | None |
| P422A | PV | 20 | male | DNMT3A Pro904Leu 18% |
| P427A | PV | 44 | female | None |
| P443B | PV | 15 | male | None |
| P483C | PV | 76 | female | None |
| P486B | PV | 52 | male | NFE2 Glu297_Arg300del 15% |
| P489A | PV | 8 | male | None |
| P490B | PV | 38 | male | TERT Ala801Thr 25% |
| P497A | PV | 80 | male | None |
| P498A | PV | 63 | male | None |
| P500A | PV | 70 | female | None |
| P506A | PV | 100 | male | None |
| P508B | PV | 83 | male | None |
| P520A | PV | 83 | female | None |
| P529A | PV | 36 | male | ASXL1 Val515fs* 37%; TET2 Asn170fs* 8% |
| P532A | PV | 45 | male | None |
| P545A | PV | 31 | male | Not studied |
| P559A | PV | 51 | male | Not studied |
| P288A | ET | 31 | female | None |
| P317A | ET | 7 | female | None |
| P356A | ET | 19 | male | None |
| P365A | ET | 25 | female | None |
| P368A | ET | 2 | female | None |
| P369A | ET | 8 | female | None |
| P379A | ET | 25 | male | None |
| P414A | ET | 35 | female | None |
| P423A | ET | 7 | female | None |
| P432A | ET | 1 | female | None |
| P433A | ET | 13 | female | None |
| P437A | ET | 29 | male | None |
| P442A | ET | 9 | male | None |
| P460A | ET | 17 | male | None |
| P485A | ET | 16 | female | None |
| P501A | ET | 3 | female | None |
| P507A | ET | 100 | male | TET2 Ser585X 46%; TET2 Phe785fs* 46% |
| P510A | ET | 28 | female | None |
| P536A | ET | 9 | male | Not studied |
| P537A | ET | 15 | female | Not studied |
| P544A | ET | 33 | male | Not studied |
| P560A | ET | 3 | male | Not studied |
| P203A | PMF | 87 | male | None |
| P212A | PMF | 47 | male | None |
| P220A | PMF | 37 | male | ASXL1 Gly646Trpfs* 30% |
| P253A | PMF | 51 | male | CBL Lys382Arg 34%; SF3B1 Lys666Arg 19% |
| P300A | PMF | 40 | male | None |
| P316A | PMF | 10 | male | None |
| P322A | PMF | 45 | male | None |
| P350A | PMF | 47 | female | ASXL1 Arg965X 53% |
| P351A | PMF | 5 | female | None |
| P360A | PMF | 22 | male | CRIM1 Asn406Ser 39%; HIF3A Asp558Asn 65% |
| P370A | PMF | 35 | male | None |
| P373A | PMF | 43 | female | None |
| P376A | PMF | 51 | male | ASXL1 Asp1004fs* 41%; IDH2 Arg140Gln 43%; U2AF1 Ser34Ala 36% |
| P415C | PMF | 73 | male | None |
| P425A | PMF | 45 | male | JARID2 Ser949fs* 44% |
| P448A | PMF | 88 | female | None |
| P455A | PMF | 57 | male | ASXL1 Pro1324fs* 37% |
| P464A | PMF | 12 | male | None |
| P484A | PMF | 96 | male | None |
| P488A | PMF | 47 | male | None |
| P492A | PMF | 14 | female | None |
| P525A | PMF | 45 | male | TP53 Arg337Leu 47%; TP53 Val143Met 48% |
| P534A | PMF | 34 | female | MPL Tyr591Asp 24% |
| P548A | PMF | 59 | female | Not studied |
| P550A | PMF | 47 | female | Not studied |
| P561A | PMF | 46 | female | Not studied |
| P523A | Prefibrotic PMF | 23 | male | None |
| P527A | Post-ET MF | 80 | male | None |
| P531A | post PV-MF | 78 | male | None |
| P526B | post PV-MF | 79 | female | None |
